# Precise and scalable metagenomic profiling with sample-tailored minimizer libraries

**DOI:** 10.1101/2024.12.22.629657

**Authors:** Johan Nyström-Persson, Nishad Bapatdhar, Samik Ghosh

## Abstract

Reference-based metagenomic profiling requires large genome libraries to maximize detection and minimize false positives. However, as libraries grow, classification accuracy suffers, particularly in k-mer-based tools, as the growing overlap in genomic regions among organisms results in more high-level taxonomic assignments, blunting precision. To address this, we propose sample-tailored minimizer libraries, which improve on the minimizer-LCA (lowest common ancestor) classification algorithm from the widely used Kraken 2 [1]. In this method, an initial filtering step using a large library removes non-resemblance genomes, followed by a refined classification step using a dynamically built smaller minimizer library. This 2-step classification method shows significant performance improvements compared to the state of the art. We develop a new computational tool called Slacken, a distributed and highly scalable platform based on Apache Spark, to implement the 2-step classification method, which improves speed while keeping the cost per sample comparable to Kraken 2. Specifically, in the CAMI2 [2] “strain madness” samples, the fraction of reads classified at species level increased by 3.5x, while for *in silico* samples it increased by 2.2x. The 2-step method achieves the sensitivity of large genomic libraries and the specificity of smaller ones, unlocking the true potential of large reference libraries for metagenomic read profiling.

## Introduction

### Metagenomics and increasing library sizes

Next-generation sequencing (NGS) has revolutionized our understanding of the underlying code of living systems, offering unprecedented insights into the structure and function of genomes of different species. Reference-based metagenomics tools rely on assembled libraries of annotated genomes to classify and characterise the taxa present in a sample. While sequence alignment can be powerful but computationally complex, k-mer based tools offer an attractive cost/performance tradeoff. Kraken [3] introduced the concept of metagenomic classification based on exact matching against a database of k-mers and lowest common ancestor (LCA) based taxonomic assignment. This allows for very rapid classification of reads. Kraken 2 [1] refined this concept to store only minimizers - minimal subsequences by some ordering - of k-mers together with their LCA taxa, yielding similar performance with much smaller databases. Kraken 2 also employs a spaced seed mask to tolerate nucleotide variance within minimizer strings, improving recall. It also uses a compact hash table (CHT) for greater efficiency, though this can lead to spurious matches when unrelated minimizers share a hash. By using Bracken [4] together with Kraken 2, database bias stemming from the selection of reference genomes can also be removed, which also allows the user to estimate the true read abundance for each taxon (read profiling).

The accuracy of reference-based metagenomics relies on the comprehensiveness of the reference libraries. Ideally, the reference library should include all the true genomes present in the sample to ensure accurate and reliable classifications. In recent decades, significant efforts have expanded many publicly available, highquality annotated genome libraries. Since its first release in 2003, the NCBI RefSeq genome repository has grown rapidly, with the number of bacterial genomes doubling every 1.5 years [5].

Larger and more comprehensive reference libraries are essential for accurate classification, as current research emphasises their role in reducing erroneous classifications [6, 7, 8]. Even when focusing on a specific taxonomic group (e.g., fungi, viruses, bacteria), false positive rates can remain high if the reference database is limited to only that group [7]. Moreover, genome libraries are often noisy, plagued by sequence contamination and taxonomic misannotations [6]. Contaminants from kits or lab processes can also lead to misclassifications, particularly when using highly specific or incomplete libraries that do not account for sequencing artefacts or unrepresented genomes [9]. In general, the selection and curation of reference genomes is an essential step, which the Kraken 2 and Bracken authors also highlight [10].

### 2-step classification

Unfortunately, increasing library size can reduce classification specificity. As libraries expand, each read has more possible genome matches, complicating precise classification. This issue is pronounced with tools like Kraken 2, which assigns shared k-mers to the lowest common ancestor (LCA) of matching taxa. With larger libraries, LCAs tend to shift up the taxonomic tree, resulting in more classifications at higher taxonomic ranks [5]. This makes classifications more *vague*, impacting species-level accuracy (*specificity*), as shown in fig. 1. Notably, the number of species in the NCBI RefSeq library has been doubling faster than the number of genera [5], suggesting that larger libraries disproportionately hinder species-level classification. In theory, longer k-mers and minimizers can increase the maximum number of distinct records in the minimizer database as the library grows, solving the problem of elevated LCAs. However, in the case of conserved regions between different reference genomes, such longer minimizers may not help, as the longer sequences would still be shared between genomes, keeping LCA taxa at a higher rank. In addition, longer minimizers may also lose recall since they are more sensitive to nucleotide variance.

**Fig. 1:**
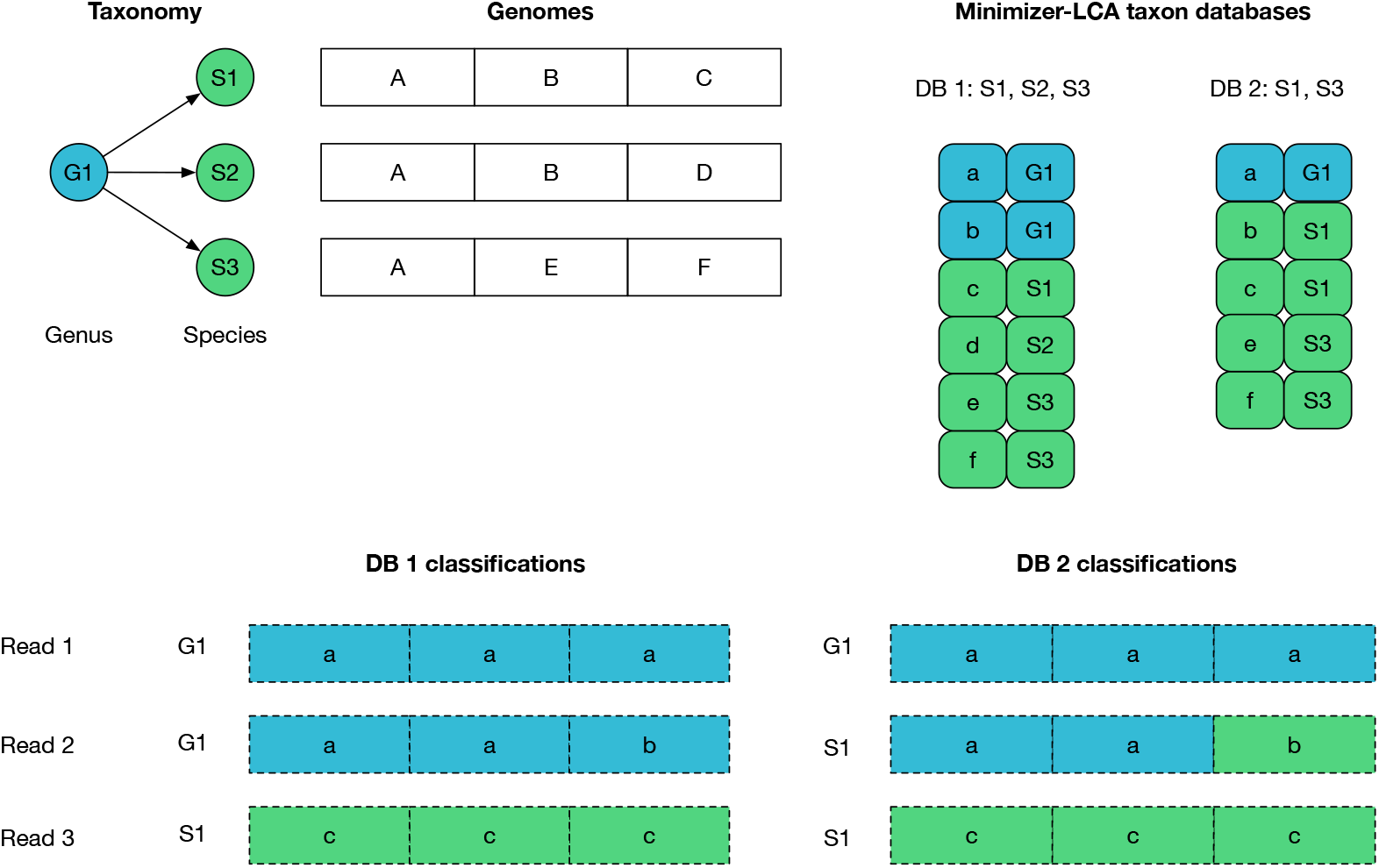
Excluding genomes when the minimizer database is built produces different classifications for reads from other genomes. In this toy example, a genus G1 contains three genomes S1, S2, and S3. These genomes contain the regions *A*..*F* which contain the minimizers *a*..*f*. Some regions are conserved across genomes. Database 1 contains all three genomes which leads to six minimizer records, two of which are on genus level and four on species level. In database 2, the genome S2 has been removed, which pushes minimizer *b* down to species level. Read classification is by majority vote as in Kraken 2: the highest weighted path from leaf taxon to root is the final classification of a read. Database 1 classifies two reads as G1 and one read as S1, whereas database 2 pushes read 2 down to species level.

Some tools alleviate problems arising from large reference libraries by removing non-resemblance genomes from the library in an intermediate step [11]. However, to the best of our knowledge, this method has not yet been applied in the context of k-mer based read binning.

We propose a novel *2-step classification method* that tailors the reference library to each sample. First, a preliminary classification with a large library conservatively filters out genomes not present in the sample. Next, a dynamic LCA-minimizer database containing only those genomes deemed to be present is built, and a second, more precise classification is performed. This approach leverages knowledge of the entire sample metagenome to improve per-read classification specificity. Using a maximally large library for the first step reduces the risk of missing a genome, which could lead to false negatives and even false positives, while using a small but highly specific library, *built specifically for the samples being classified*, for the second step alleviates the vagueness problem.

### Slacken

Directly implementing the 2-step method with Kraken 2 is computationally costly due to the time required to dynamically build sample-tailored databases. Additionally, Kraken 2 can require highly specialised computing resources with terabytes of RAM, and running it on large genome libraries—currently considered best practice for most reliable and accurate results [8] —is already challenging. Instead, to study the 2-step method, we developed a new implementation of the Kraken 2 algorithm designed to run on modern scalable infrastructure, called Slacken. Slacken implements the original Kraken 2 algorithm as faithfully as possible (1-step method), while also building large databases more efficiently. Slacken also implements the 2-step classification method efficiently, allowing us to study this method in a variety of settings.

Slacken is built using Apache Spark (https://spark.apache.org) [12], a highly scalable distributed computing framework, and runs both on single machines and on clusters. It is compatible with Kraken 2, in the sense that it accepts the same input file formats and produces compatible output file formats. The Fastdoop [13] library is used for efficient sequence input. Unlike Kraken 2, Slacken does not need to keep the entire database in memory, which allows for scaling to databases larger than the total RAM. Crucially, Slacken can classify multiple metagenomic samples at the same time, creating a large multisample read pool prior to classification and grouping back reads by sample after. This helps reduce the per-sample cost of classification, especially when using large genomic libraries.

In 2-step classification, multisample classification is a performance benefit, as in the 1-step method, but also influences the contents of the second library: including additional samples may cause additional taxa to be detected in the first step, which lets their genomes be included as possible classification targets in the second step. Thus, every sample can influence the classification results of every other sample. For this reason we classify related samples together.

Figure 2 contrasts the dataflow in 2-step Slacken (described below) with regular (1-step) Kraken 2 classification. Kraken 2 relies on a pre-built static LCA-minimizer database for classifications. Slacken uses a static database for the initial classification (the first step) only. In the second step, a heuristic is used to select taxa from the results and a new, smaller (in our experiments, typically around 10% or less of the initial size by minimizer count) LCA-minimizer database is built, containing only minimizers from the detected taxa. This is equivalent to building a second Kraken 2 database on the fly with `kraken2-build`. All reads are now classified again using this second database, which produces the final result. Slacken’s operations are thus equivalent to `kraken2` (classify), `kraken2-build`, and (optionally) `bracken-build`, with the important difference that multiple samples can be classified simultaneously. As with Kraken 2, the final output is a read binning and a read profile (report), and Bracken may be used to remove database bias from the latter, accurately estimating the true read abundance for each taxon in the sample.

**Fig. 2:**
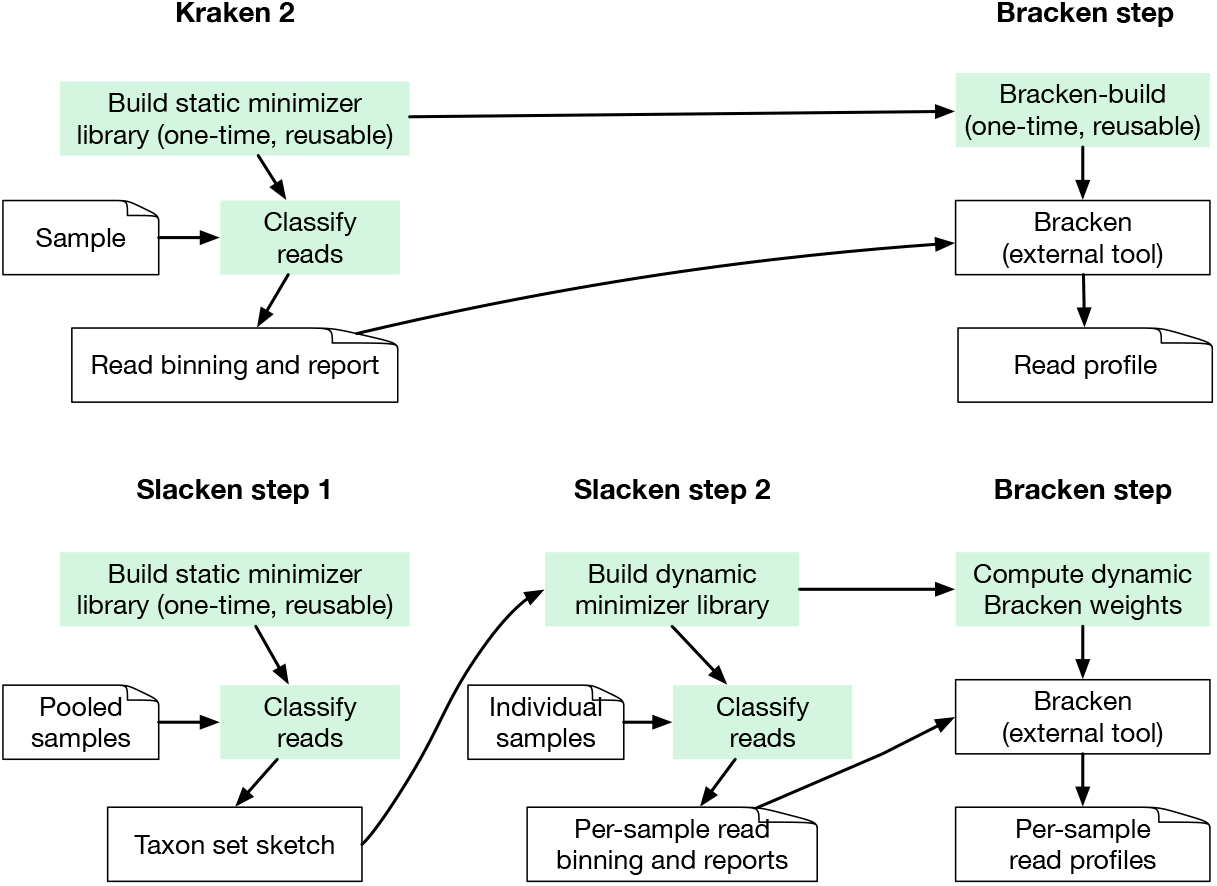
Kraken 2 + Bracken (top) compared with Slacken 2-step + Bracken (bottom) internal data flow. Coloured boxes are equivalent between the two systems in the sense that they compute the same results. Slacken and Kraken 2 both build a reusable minimizer-LCA taxon library which is the basis for classification. However, Slacken optionally pools all samples together for the first step. Pooled classification results in a taxon set sketch, which is the basis for a second minimizer-LCA library that is built on the fly (dynamic). Samples are then classified again using this second library. Bracken weights are also computed on the fly from the dynamic library, which means that Bracken may be run directly on the outputs from Slacken.

## Materials & Methods

We evaluate the taxonomic read-binning performance of Slacken against Kraken 2, as well as the downstream taxonomic profiling results of Slacken + Bracken against Kraken 2 + Bracken.

Experiments were run on Amazon Web Services EC2 instances. Slacken was run on AWS Elastic MapReduce (EMR), on instances from the `m6gd` and `m7gd` families, which have 4 GB of RAM per CPU and physical NVMe drives. We used the `4xlarge` (16 CPU) and `8xlarge` (32 CPU) instance sizes. Genomes, samples, and built minimizer databases were stored on Amazon S3. AWS EMR release 7.1.0 was used, which includes Spark 3.5.1 and Java 17.

Kraken 2 version 2.1.3 and Bracken version 2.9 were used. These were run on instances of the type `x2gd.metal`. Kraken 2 was run in memory-mapped mode with the databases pre-copied to `/dev/shm` (shared memory) for best performance. During Slacken development, when we initially validated its correctness, we used a custom built version of Kraken 2 with debug code inserted, to be able to trace the origin of each minimizer record. For all other experiments, however, the unmodified version of Kraken 2 was used.

For all experiments, the default Kraken 2 parameters of *k* = 35, *m* = 31, *s* = 7, as well as the default spaced seed mask and XOR mask were used for both Kraken 2 and Slacken.

### Two-step classification

We investigated the impact of applying a read based cutoff (R) filter for genome inclusion into the 2-step library. More specifically, given a read cutoff of N reads, we include in the dynamic library the genome of a species level taxon T and the genomes of all its taxonomic children if the total number of reads assigned to T and its children is at least N in the first classification. In all of our experiments we used multi-sample classification, which means that the number of reads for a given taxon needed to be at least N in total across all the samples in a group pooled together. We compare results for *R* ∈ {1, 10, 100}.

Details on the taxon set discovered by each heuristic for the various genomic libraries and sample families are available in supplementary table S3.

### Genomic libraries

Two different genome libraries were used to build databases for this study. Both libraries are subsets of RefSeq release 224.

We used the Kraken 2 build scripts to obtain and preprocess genomes. The same low-complexity masking as for Kraken 2 was used (`k2mask`).

- **Standard (std)**: The selection of genomes included in the Kraken 2 standard library. This includes complete genomes and chromosomes from archaea, viruses, bacteria, plasmids, and human, as well as adapter sequences (UniVec core). This was used to build databases with both Slacken and Kraken 2.
- **RefSeq prefer complete (rspc)**: The NCBI Reference Sequence genome library with only complete genome sequences/chromosomes for those taxa that have them, and draft genomes/scaffolds otherwise, as well as adapter sequences (UniVec core). This allowed us to cover as many taxa in RefSeq as possible, while avoiding contamination and noise from incomplete genomes for those taxa that had a complete genome. This was used to build databases with Slacken only.

Here, the std library is a strict subset of the rspc library. Both libraries include genomes from archaea, bacteria, human, viruses, plasmids, as well as adapter sequences. In addition, rspc contains fungi and other eukaryotic genomes.

These two libraries are compared in table 1. The number of taxa here denotes taxa with associated genomic sequences; the number of nodes in the complete tree when parent taxa are included is larger.

**Table 1.**
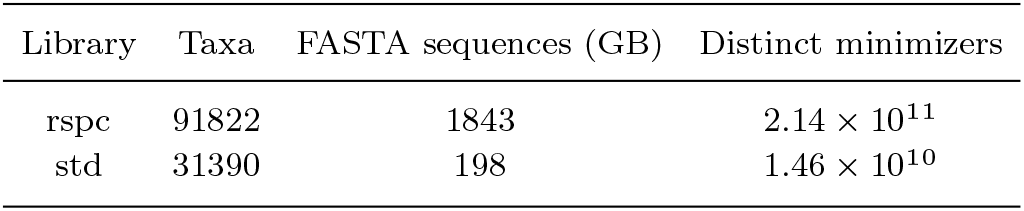
Comparison of minimizer-LCA libraries.

| Library | Taxa | FASTA sequences (GB) | Distinct minimizers |
| --- | --- | --- | --- |
| rspc | 91822 | 1843 | $2.14 \times 10^{11}$ |
| std | 31390 | 198 | $1.46 \times 10^{10}$ |

### Classification experiments

Slacken and Kraken 2 were used to classify a total of 160 samples (see below). A total of 7040 sample classifications were compared against the ground truth reference at both the species and genus levels, for a total of 14080 experiments. Detailed results for each experiment are available in supplementary table S1, and visualizations are available in the supplementary figures.

We evaluated both Kraken 2 and Slacken classifiers at classification confidence levels of 0, 0.05, 0.1 and 0.15. An exhaustive analysis of Kraken2’s parameter space by [8] identifies 0.15 as an optimal confidence threshold, which we primarily used to measure classifier performance although similar trends hold at other confidence levels. We used Kraken 2 and 10 Slacken-based classifiers for performance comparison and to investigate the impact of different values for the read based cutoff filter:

- **rspc R100/std R100**: Slacken with 2-step classification and a preliminary classification with a rspc/std database, 1-step confidence (c) at 0.15 and 2-step taxon filter at ≥ 100 reads per taxon.
- **rspc R10/std R10**: Slacken with 2-step classification and a preliminary classification with a rspc/std database, 1-step confidence (c) at 0.15 and 2-step taxon filter at ≥ 10 reads per taxon.
- **rspc R1/std R1**: Slacken with 2-step classification and preliminary classification with a rspc/std database (c) at 0.15 and 2-step taxon filter at ≥ 1 reads per taxon.
- **rspc gold/std gold**: Slacken with 2-step classification where the 2-step database is built using genomes from ground truth taxa and their descendants in rspc/std.
- **rspc 1-step/std 1-step**: Slacken 1-step classification with rspc/std database.
- **kraken2**: Kraken 2 classification with the std database.

For gold set classifiers, the gold set was computed as the union of ground truth taxa assigned to all reads of all samples in the corresponding sample family.

### MetaPhlAn Analysis

To compare taxon set precision and recall statistics for Slacken against the state of the art in taxon profiling, i.e., MetaPhlAn [14], we classified all 160 samples (see below) using MetaPhlAn 4.1.1 with the marker database ChocoPhlAn version `mpa vJun23 CHOCOPhlAnSGB 202403`. Profiling was done using the flag `--add viruses`. Both were the latest available versions of the tool and database at the time of writing.

We considered all taxa with nonzero abundance in the MetaPhlAn output as detected by the profiler. For comparison with Slacken and Kraken outputs, we included all species-level classifications along with their corresponding NCBI taxon IDs, as automatically provided by the tool.

### Differences between Slacken 1-step and Kraken 2

Given the same values of *k* (k-mer width), *m* (minimizer width) and *s* (spaced seed mask spaces), and given the same genomic library and taxonomy, Slacken classifies reads using the same algorithm as Kraken 2. However, Slacken uses a different internal data structure, as shown in Figure 3. Kraken 2 hashes minimizers, which are furthermore stored in a compact hash table (CHT). This method is memory efficient but introduces potential collisions between unrelated minimizers, which can randomly lift LCA taxa to a higher rank. We refer readers to [1] for details on the collision rate of the CHT. Slacken does not hash minimizers but instead stores them in full. This means that each minimizer record in Kraken 2 corresponds to at least one record in Slacken, and several in the case of collisions.

**Fig. 3:**
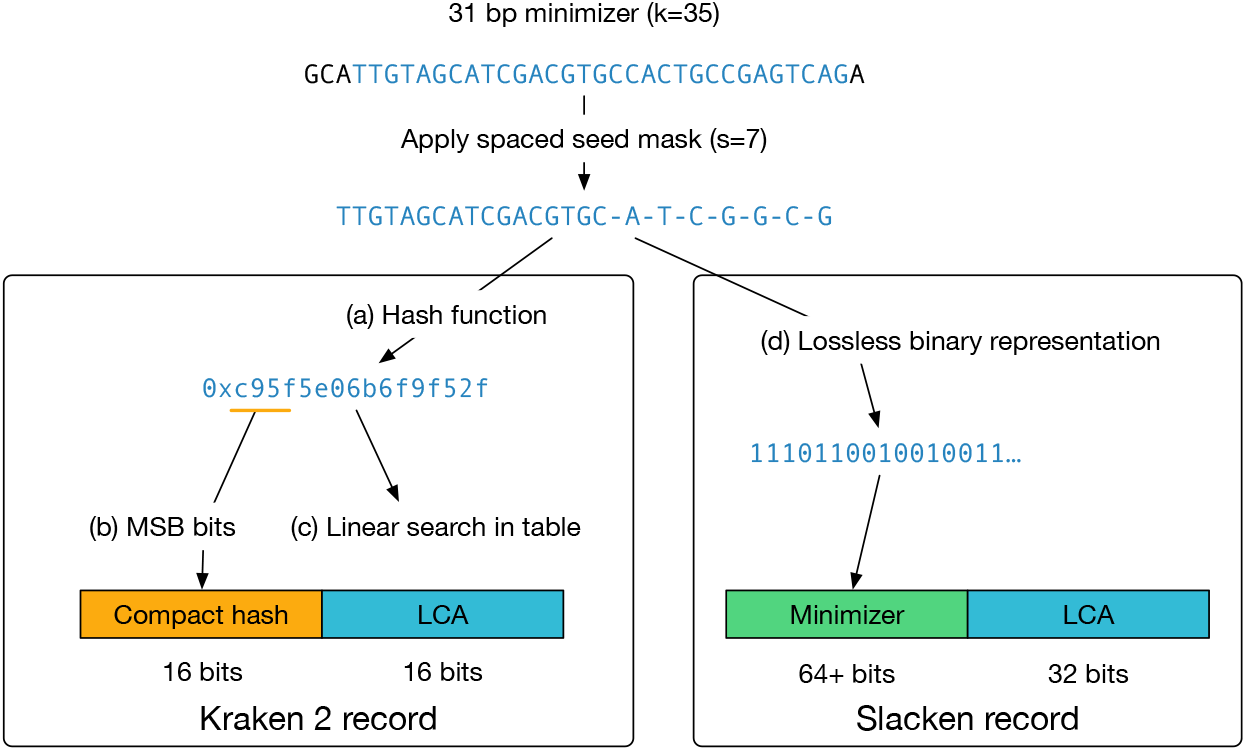
The minimizer-LCA records used in Kraken 2 and Slacken. Both tools start from an *l*-length minimizer in a *k*-length window (31 and 35 respectively, with the standard parameters) and apply a spaced seed mask (7 spaces by default). Kraken 2 then applies a hash function (a), from which the most significant bits become the compact hash code (b) and the entire hash code is used to perform a linear search in a compact hash table (c). These three steps potentially introduce collisions between biologically unrelated records. Slacken does not hash the minimizer but instead encodes the minimizer using two bits per nucleotide (d). The number of compact hash bits in Kraken 2 shrinks when the number of LCA bits grows, as one record must fit in 32 bits. For the standard database using RefSeq 224, this was 16 hash bits. Slacken uses as many minimizer bits as needed (2*l*, up to 256) and 32 bits per LCA taxon.

### Statistical analyses

All simulated and mock datasets contained per-read taxon labels. For each classified read, following the methodology used by Wright [8], we assessed it as

- **True positive (TP)**: If the true label is identical to, or an ancestor of the classified label, or if the classified and true labels share the same species-level ancestor. For example, if a the true label is *Streptococcus agalactiae* and the classified label is the descendant strain *Streptococcus agalactiae 18RS21*.
- **Vague positive (VP)**: If the true label is a descendant of the classified label and the classified label is above species level. For example, if the true label is *Bacteroides fragilis* and the classified label is *Bacteroides*.
- **False positive (FP)**: If the true label is neither a descendant nor an ancestor of the classified label. For example, if the true label is *Bacteroides fragilis* and the classified label is *Streptococcus*.
- **False Negative (FN)**: If the classifier failed to assign a label to the read.

Like Kraken 2, Slacken classifies reads at multiple levels in a taxonomic hierarchy. We define true positives (TP) and vague positives (VP) relative to the species level. Given the limited strain-level resolution in genomic libraries relative to the vast number of existing strains, we consider classification at this level sufficiently precise. Consequently, if a read is classified to the correct species-level ancestor of a given strain, we count it as a TP.

We also computed the following metrics for measuring the quality of the detected taxon set at the species level. Here, TP, FP and FN refer to counts of detected taxa at the species level:

- **Taxon Set Precision**: precision of species level taxa detections: 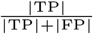
- **Taxon Set Recall**: recall of species level taxa detections: 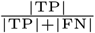

### Index metrics

We use two variants of the index metric as introduced in [8].

We assign to each classified read an integer between 0 and 9, called the read-index, as follows:

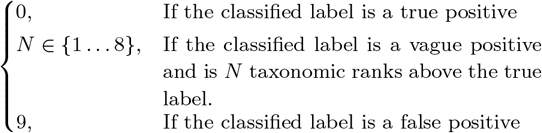

The number of ranks in the taxonomic tree is 8, which is why FPs are given a weight of 9.

We define two types of indexes:

- **Soft read index**: The soft index for a read (or s-index) is computed using rules 1-3 along with the additional rule that false negative read classifications (i.e. unclassified reads) are assigned an index 0.
- **Read index**: The index for a read is computed using rules 1-3 along with the additional rule that false negative read classifications (i.e. unclassified reads) are assigned an index 9.

The average read index (or s-index) for all reads in a sample is called the *sample index*, or *sample s-index*, respectively). Both indexes can be defined for a given taxonomic rank by promoting all true positive labels up to that given rank and then computing the index as above. Figures 4 and 8 show index and s-index values for samples at species ranks.

**Fig. 4:**
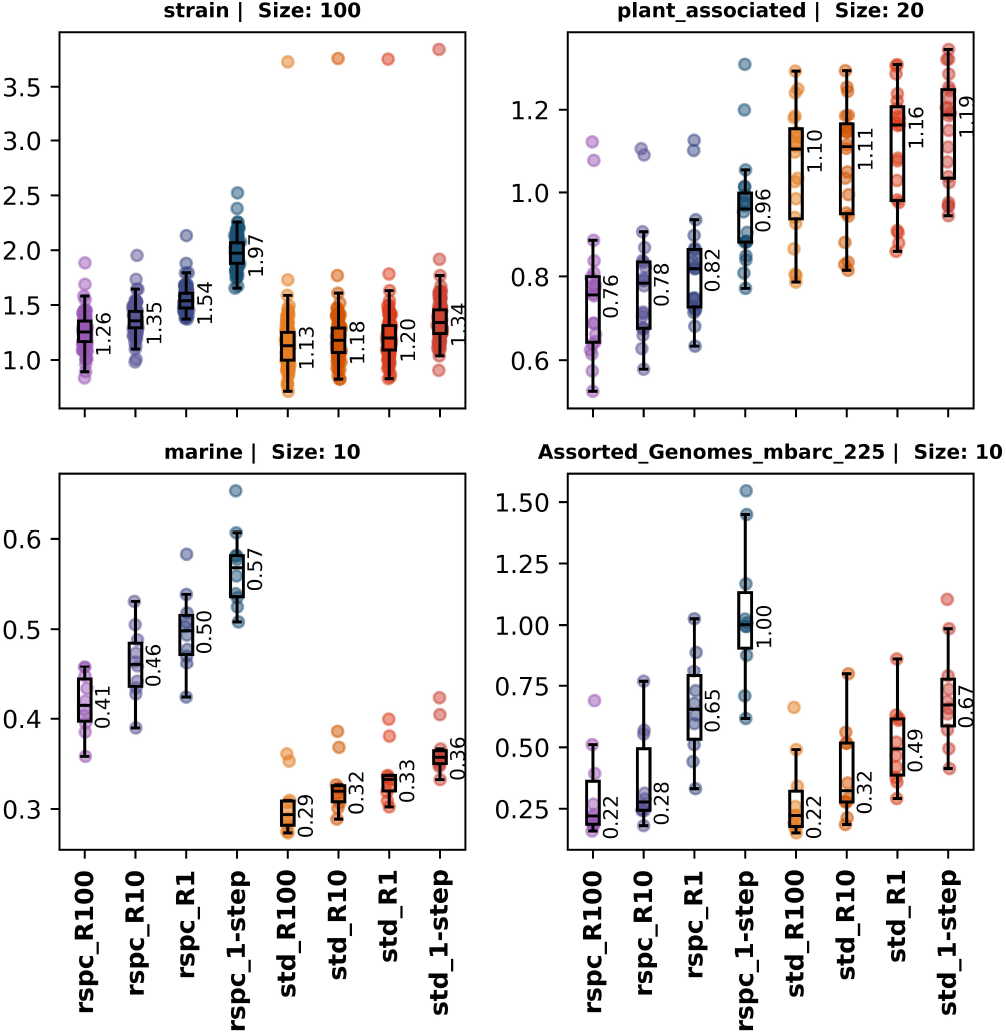
Boxplots of sample soft-index values for strain, plant associated, marine and mbarc datasets with various 1- and 2-step classifierers. The sample soft-index is computed by taking a weighted average of all sample reads, giving a weight of 0 to TPs and FNs and 9 to FPs. VP reads are given a weight corresponding to the number of ranks between the true taxon and the classified taxon label for that read.

The index metrics defined here deviate from their original conception in [8]. Unlike the original, which assigns an *fp* read an index equal to the number of ranks to the common ancestor of the *fp* taxon label and the ground truth, our approach simplifies comparability and interpretation. By design, a lower index in our framework directly reflects higher classification precision.

### Bracken experiments

Each of the 7040 classifications was also further processed with Bracken to compensate for library bias. Post-Bracken results for each experiment are available in supplementary table S2 and visualisations in the supplementary figures.

The ground truth was for the CAMI2 samples obtained from the CAMI2 gold standard mapping, where reads that had no species-level mapping were first filtered out. Since Bracken by default redistributes every read to species-level, this allowed us to compare with a true read profile for each sample. For *in silico* samples, the ground truth was known by construction. The default Bracken cutoff of 10 reads for a taxon to be present was used in all cases.

Each sample was represented as a vector with entries in the range [0, 1] where each entry represented the fraction of reads assigned to that taxon. We measured the following metrics:

- **LSE**: Euclidean distance from the ground truth profile: LSE(*x, t*)
- **LSE log-10**: LSE(log(*x*), log(*t*))
- **L1**: L1 (Manhattan) distance from the ground truth mapping: L1(*x, t*)
- **L1 log-10**: L1(log(*x*), log(*t*))

### Simulated and mock sample dataset characteristics

We evaluated 2-step and 1-step classification across 6 sample families, of which 3 were simulated and 3 were mock sample datasets.

- **Simulated Communities**: Short-read samples simulated for the marine (number of samples (*n*): 10) and rizosphere (plant associated) (*n* = 20, samples 0-19), microbial environments as well as the strain madness (*n* = 100) dataset from the well known CAMI2 challenges [2]. These datasets contain simulated reads from a combination of reference and novel genomes, assembled from real metagenomic datasets and are widely used for benchmarking metagenomics tools, including classifiers and binners. Each sample contained, after filtering, on average 7.76 × 10^6^ read pairs.
- **Mock Communities**: Short-read samples simulated from 225 complete genomes chosen at random from Kraken 2’s std database. Assorted Genomes 225 (*n* = 10) and Assorted Genomes mbarc 225 (*n* = 10) are simulated using CAMISIM [15] with the former simulated using the less noisy Illumina HiSeq quality profile from ART [16] and the latter, using the quality profile used to simulate the three CAMI2 datasets, i.e. the mbarc profile. The third dataset, i.e. Assorted Genomes Perfect 225 (*n* = 10) was simulated in an error free manner using CAMISIM. Each sample contained on average 3.46 × 10^6^ read pairs.

The CAMI2 datasets were generated using genome contigs assembled from real metagenomic data. We further filtered each simulated metagenome to remove reads came from genomes that had taxon labels above a species level. This was done to identify each read with a single species of origin, since the fact that a few genomes had genus or above taxon labels was a limitation of the metagenomic assembler used to generate the CAMI2 datasets and not reflective of the underlying metagenomic sample. This also made counting the true positive classifications at a given taxonomic rank unambiguous, and allowed us to measure the quality of post-Bracken read profiles directly against the reference. The amount of reads filtered out varied between on average 10% (strain madness samples) to on average 20%(plant associated samples).

The strain, marine, and plant associated datasets were selected because each presents unique challenges for metagenomic classification using the chosen genome libraries. As shown in table 2, these datasets progressively include more taxa that fall outside the std and rspc libraries, enabling us to assess whether certain trends hold consistently across datasets from varied environments, thus reinforcing the robustness of our results. The three mock datasets were created primarily to validate Slacken and serve as a stress test for classification with large libraries.

**Table 2.** Number of distinct taxa and average reads per sample in datasets.

| Dataset | In rspc | In std | In cfl<sup>1</sup> | Total | Reads (M) |
| --- | --- | --- | --- | --- | --- |
| Strain | 19 (1) | 18 (2) | 17 (3) | 20 | 6.13 |
| Marine | 465 (34) | 448 (41) | 404 (85) | 489 | 14.10 |
| Plant | 261 (50) | 156 (155) | 225 (86) | 311 | 12.70 |
| A.G<sup>2</sup> | 197 (1) | 198 | 114 (84) | 198 | 3.33 |
| A.G.mbarc | 197 (1) | 198 | 114 (84) | 198 | 3.33 |
| A.G.Perfect | 197 (1) | 198 | 114 (84) | 198 | 3.70 |

<sup>1</sup> *ChocoPhlAn*, the MetaPhlAn marker database used
<sup>2</sup> *Assorted\_Genomes*

Number of taxa from the sample families present in each library, as well as the average number of reads (read pairs, in millions) per sample. The number of novel taxa, if any, is given in parentheses. In the rspc library, all minimizers from *Haloferax alexandrinus* had been lifted to genus level, effectively giving the A.G. samples one novel genome.

Similar to the CAMI2 strain dataset, but simulated directly on genomes present in std and rspc, the results allowed us to show that Slacken 2-step classification using rspc can still recover comparable precision to Kraken 2 on std. This is a pathological case where using a larger library is a clear disadvantage.

### Speed and cost of classification

Speed and cost were evaluated on 50 samples (IDs 0 to 49) from the strain dataset on AWS EMR. These were on average 4 GB each as uncompressed fastq files. Slacken was run on a 50/50 mix of regular and interruptible (spot) instances with 4 GB RAM per CPU, with a total of 512 CPUs. Kraken 2 was run on a single machine with 64 vCPUs and 1 TB RAM in memory-mapped mode. For cost measurements, the total cost of running a Slacken job was divided by the number of samples in the job to give the average cost per sample.

Due to the high cost of building a large database with Kraken 2, we are comparing rspc performance with that of a pre-built *nt* database, which was of comparable size (1.5 TB of sequence data for nt, vs 1.8 TB for rspc). As Kraken 2 performance is mainly dependent on the database size for a given sample, this approximates the cost of running classification with the rspc database.

When comparing the 2-step method with Kraken 2 for performance evaluation purposes, we simulated 2-step classification as follows. We used `kraken2` to first classify once (estimating the cost using the nt database, as above), then using `kraken2-build` to build a custom database based on the taxon set that would have been detected (using the rspc genomes), classifying with this second database using `kraken2`, and when applicable, running `bracken-build` on the result to generate Bracken’s k-mer distribution.

## Results

### Slacken 1-step implements the Kraken 2 method faithfully

To establish Slacken’s overall fidelity to Kraken 2, for each sample studied in the present work, we measured the L1 distance between the classification report (fraction of reads assigned per taxon) generated by 1-step Slacken and the one generated by Kraken 2, using the std library. The distance was never larger than 0.005, corresponding to 0.5% of reads.

During validation, we found that Kraken 2 during its build process finds spurious minimizers immediately after ambiguous regions in genomes, seemingly incorrectly. While this is not harmful, it gives Kraken 2 a slightly larger minimizer database for the same parameters. For the Kraken 2 standard library, we found that the difference due to this issue was about 1% of the total minimizer count.

### 2-step classification recovers specificity loss in growing reference libraries

We used three simulated and three in-silico datasets to compare the taxonomic binning performance of Slacken 1-(or Kraken 2) and 2-step methods, as well as their effectiveness as taxonomic profilers when used with Bracken. The simulated datasets—strain madness (strain), marine, and plant associated—were sourced from the CAMI2 challenge [2] and are based on genomes assembled from actual sample metagenomes. The in-silico datasets were custom-designed and based on genomes from the standard Kraken2 library. We used the first three datasets to assess performance in a realistic scenario with unknown or novel genomes, while the latter three allowed us to evaluate classifier performance in an idealized controlled environment with only simulated sequencing noise. We present results below using the Assorted Genomes mbarc 225 (mbarc) in-silico dataset as the representative dataset (as trends were consistent across all three in-silico datasets), as mbarc has the same noise profile used in creating the CAMI2 samples.

Large reference libraries are known to affect the accuracy of k-mer-based LCA methods [5]. We confirm this loss of classification specificity on our simulated and in-silico metagenomic datasets using both the rspc and std libraries. Figure 5 (right, unshaded region in each subplot) illustrates this effect, showing how true positive (TP) classifications with Slacken 1-step on the smaller std library become vague positive (VP) classifications, i.e. classified as some ancestor of their true taxon, with Slacken 1-step on the much larger rspc library.

**Fig. 5:**
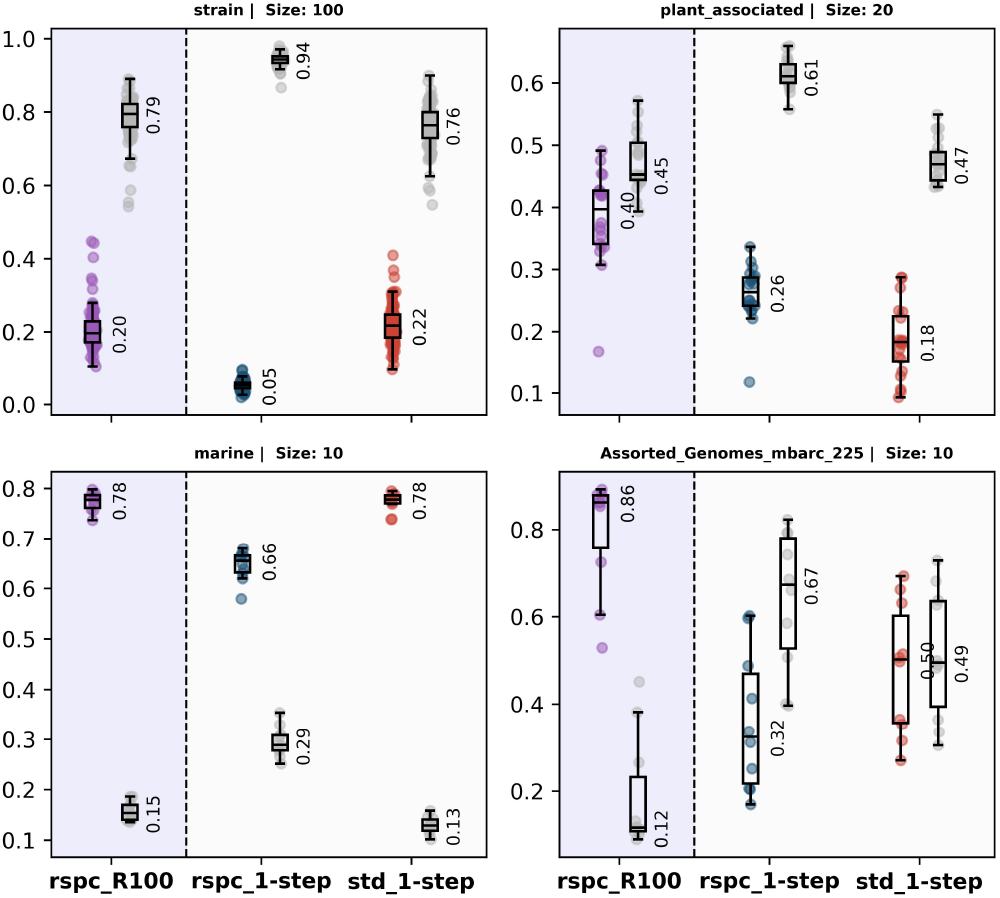
Each of the four plots contain fraction of total reads present at species-level for true positive (colour) and vague positive (grey) for the Slacken 1-step classifiers on the *std* and *rspc* library databases (right-unshaded region in each subplot). The blunting of classifications when using the larger *rspc* library is clearly shown by the decrease in TPs and increase in VPs. Median values are given beside each whisker plot.

We created eight Slacken classifiers, one for each combination of *R* ∈ {1, 10, 100} (read count per taxon cutoffs) and library choice of std or rspc, as well as two 1-step classifiers. Figure 5 (left, shaded region in each subplot) demonstrates that the Slacken classifier with *R* = 100 on rspc (rspc R100) mitigates the specificity loss due to the larger library, and in the case of the in-silico samples even improves on the initial results.

Table 3 shows the size of the 2-step library for each classifier and sample family. The improvement in specificity was possible due to the significantly smaller size of the 2-step libraries relative to the reference libraries. As expected, the 2-step library sizes decrease with higher values of *R*. For each cutoff (*R*) value we detected more taxa in the larger genomic library (rspc). This is likely due to the fact that rspc has more species and subspecies level taxa than std (see 2.1 on how taxa are included in the 2-step library). We noted for the in-silico samples with the *R* = 1 heuristic, that the number of taxa selected was highly sensitive to added noise.

**Table 3.** Number of species level taxa included in 2-step library.

| Dataset | Rspc |  |  | Std |  |  |
| --- | --- | --- | --- | --- | --- | --- |
|  | R1 | R10 | R100 | R1 | R10 | R100 |
| Strain | 1730<sup>1</sup> | 590 | 256 | 513<sup>1</sup> | 303 | 183 |
| Marine | 2253 | 1511 | 914 | 1594 | 1046 | 757 |
| Plant.Associated | 2396 | 1296 | 737 | 1890 | 1052 | 652 |
| A.G | 1082 | 197 | 175 | 488 | 201 | 182 |
| A.G.mbarc | 916 | 193 | 173 | 455 | 197 | 179 |
| A.G.Perfect | 192 | 188 | 178 | 193 | 188 | 182 |

<sup>1</sup>The strain dataset was analysed in two batches comprising of 49 and 51 samples respectively (grouped by sample ID range). This figure is the average number of the species included in the 2-step library.
Slacken 1-step classification with *std* and *rspc* use libraries with 31390 and 91822 species level taxa respectively. We observed that the largest 2-step libraries (with the R1 heuristic) are nearly 17× to 40× smaller than the 1-step libraries (i.e. *std* and *rspc*) respectively.

### The 2-step method enhances classification specificity

The 2-step method enhances classification specificity by tailoring the library to the sample metagenome. By using a smaller, sample-representative library, classifications can be more precise, as long as the library adequately represents all genomes in the sample. On the other hand, it is important not to be too selective, as incomplete libraries may give rise to false positive read classifications or unclassified reads. This limitation is particularly relevant for real samples, which often contain novel genomes.

We evaluated the performance of the read-based 2-step classifiers on the strain, marine, plant associated, and mbarc datasets. Notably, ~ 60% and ~ 43% of taxa in the plant associated and marine samples, respectively, are absent in the std library, while both datasets have only about 30% of their taxa missing from rspc. The strain dataset has only one and two taxa missing from rspc and std libraries, respectively, whereas the mbarc dataset is fully contained within std (and, therefore, rspc). This allows us to assess the performance of the 2-step classifiers across different types of microbial communities and under different levels of reference library comprehensiveness. Figure 6 displays a stacked comparison of classifications using various Slacken 1- and 2-step classifiers across these datasets. We observed that Slacken’s 2-step approach enhances species-level classifications across all datasets and reference libraries. Classifications with rspc produced more significant gains in species level classifications than with the smaller std library, yielding a 15%, 14%, and 12% increase in species classifications with rspc, and a 11%, 3%, and 5% increase in species classifications with std on the strain, plant associated, and marine datasets, respectively (see supplementary table S4). This aligns with expectations, as the reduction in the size of the 2-step library is more pronounced with rspc than with std (see table 3).

**Fig. 6:**
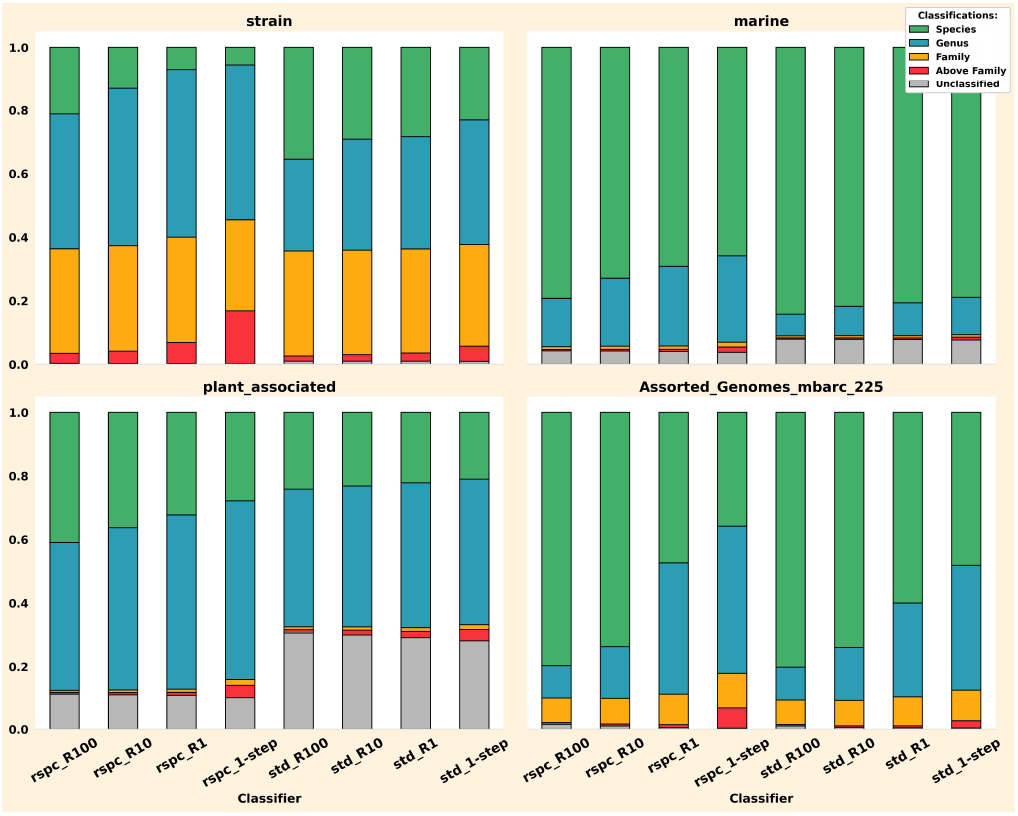
Each barplot breaks down Slacken 2-step read classifications (as a fraction of total reads) into species, genus, family, classifications above family (up to but not inclusive of root) and unclassified reads on the *std* and *rspc* libraries. 2-step classifications tend to “squash” genus, family and higher classifications and increase classifications at species ranks or leave reads unclassified.

### 2-step classification reduces overall sample index

Figure 7 shows that the increase in FPs between 1- and 2-step classifiers is negligible (~ 0.6% of total classified reads at most on an average). Unclassified reads can, however, increase by 2 − 3% (in the case of the std library), excluding outliers. This is seen in the case of the plant associated dataset which, due to the low coverage with both std and rspc has more unclassified reads. Further reducing the size of the library can increase the phylogenetic distance between novel genomes and the library which could contribute to the increased number of FNs. We therefore need a way to measure and compare the quality of 1- and 2-step classifications, taking FPs and FNs into account.

**Fig. 7:**
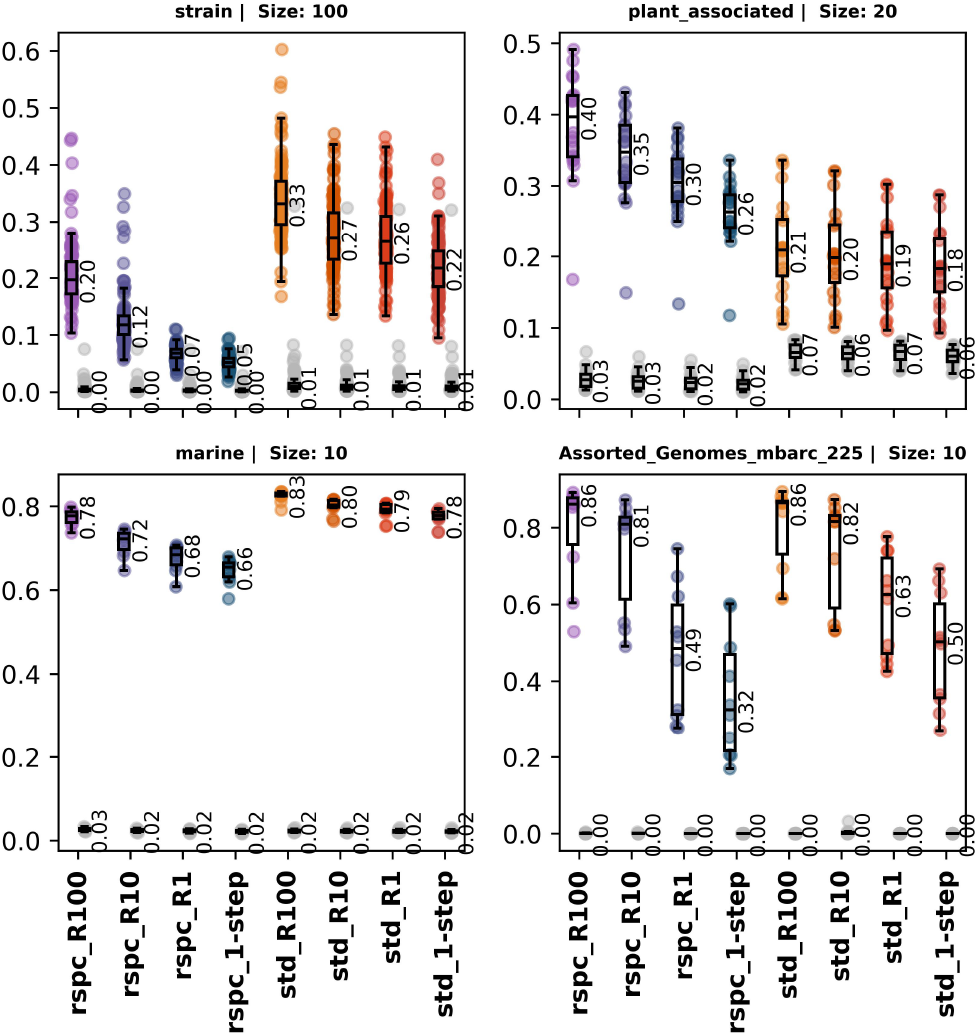
Each of the four plots contain species level read fractions of total reads present for true positive (colour) and false positive (grey) for the various 2- and 1-step classifiers on the *std* and *rspc* library databases. The increase in TP classifications is contrasted with the increase in FPs as tighter read filters are used. The increase in FPs is *<* 0.6% in both rspc and std, omitting outliers.

**Fig. 8:**
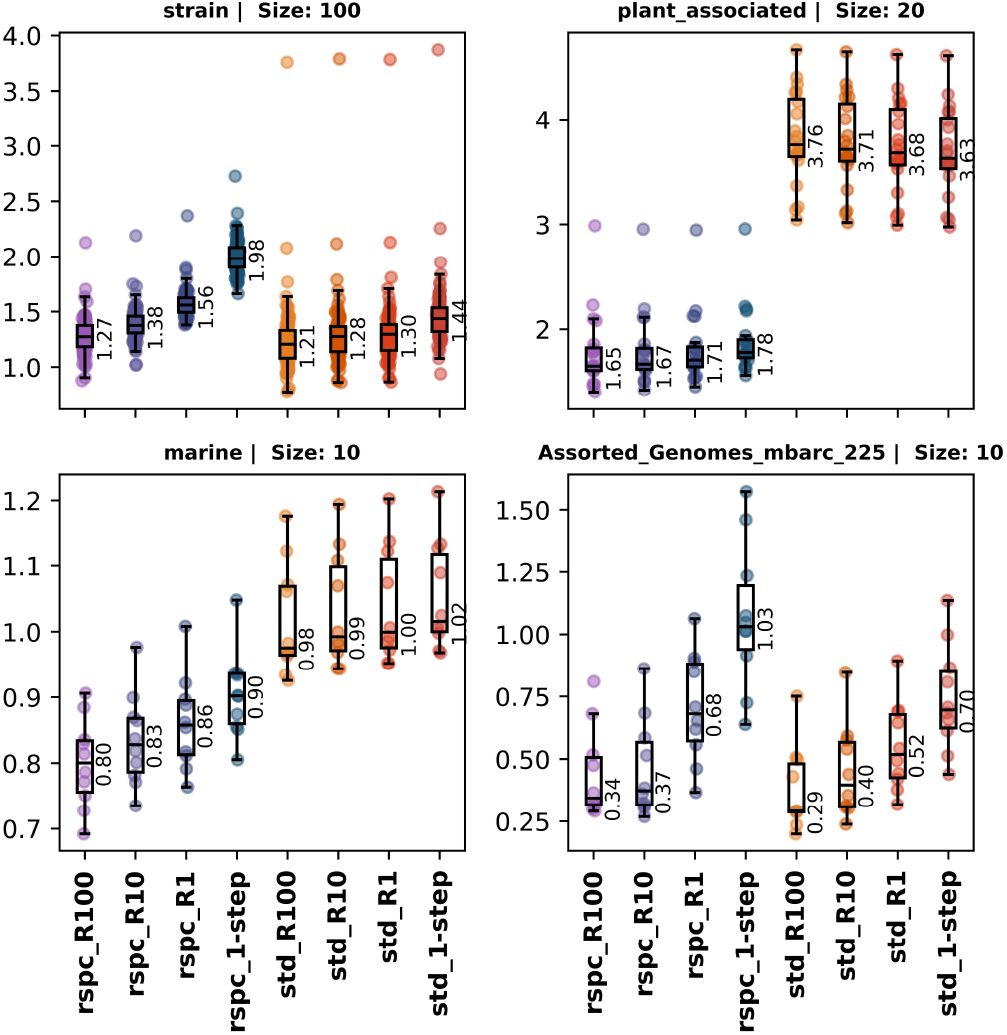
Boxplots of sample index values for strain, plant associated, marine and mbarc datasets with various 1- and 2-step classifiers. Sample index is computed by taking a weighted average of all sample reads, giving a weight of 0 to TPs and 9 to FPs and FNs. VP reads are given a weight corresponding to the number of ranks between the true taxon and the classfied taxon label for that read.

Following a concept introduced by Wright et al [8], we define the sample index as a weighted average of all classified reads within a given sample. True positive (TP) reads are assigned a weight of 0, while false positive (FP) and false negative (FN) reads receive a weight of 9. Vague positive (VP) reads are assigned a weight based on the number of ranks separating the true taxon from the classified taxon label for that read. The sample index serves as a useful comparison tool across classifications that use the same library since stricter 2-step read filters tend to yield higher FN and FP counts, although FP increases are typically insignificant. An index value of *N* implies that, on average, the classified label of a read is *N* ranks above the species level. Figure 8 shows a clear trend of decreasing index values between 1- and 2-step classifiers as the read filter is strengthened, indicating that the overall quality of read classifications improves with the 2-step classifiers.

The only exception to this trend appears in the plant associated dataset, where the std-based 2-step classifiers consistently show higher index values. This likely reflects the previously mentioned poor fit of the std library to this dataset. In this case, we analyzed the quality of read classifications while omitting unclassified reads, also called soft-index, and found that, among classified reads, the overall classification quality does indeed improve (see fig. 9).

**Fig. 9:**
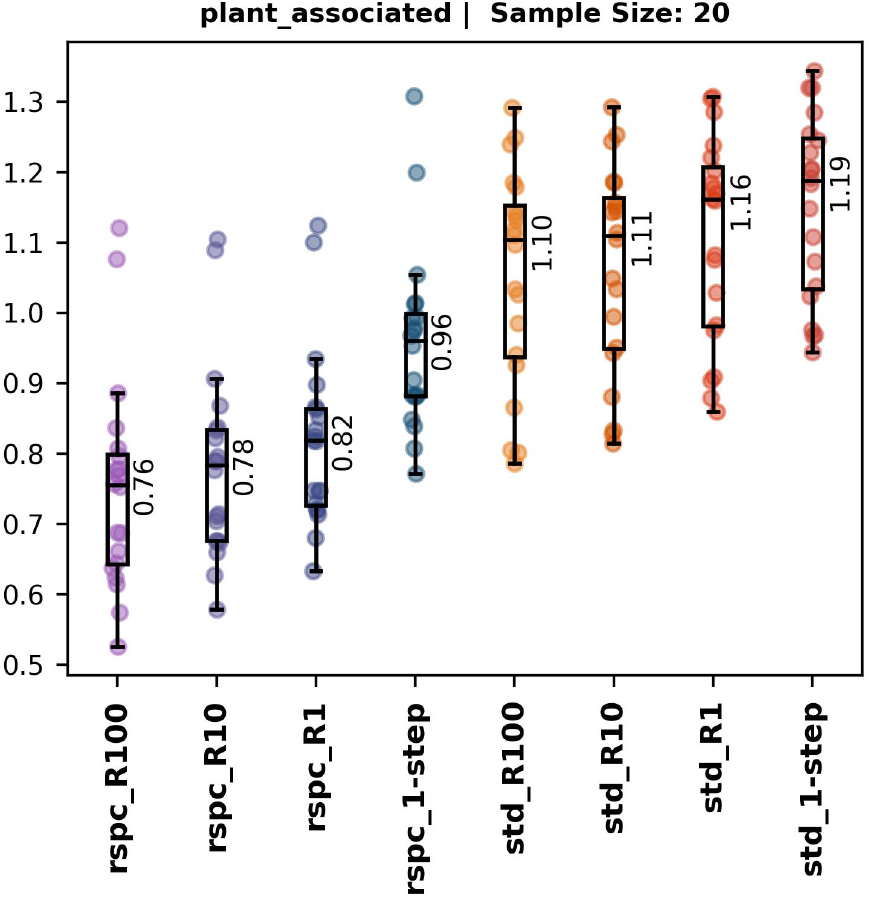
The s-index or soft index is computed by taking a weighted average of all sample reads, giving a weight of 0 to TPs and FNs, and 9 to FPs. VP reads are given a weight corresponding to the number of ranks between the true taxon and the classifieded taxon label for that read. The h-index measures the quality of read classifications amongst classified reads only.

### 2-step classification recovers lost precision in read profiling with Bracken

Slacken dynamically builds Bracken weights for 2-step libraries, enabling us to use Bracken for species-level read redistribution. We compared read fractions assigned by various 1- and 2-step Slacken classifiers against the expected ground truth profiles, using the L1 distance metric to assess the overall accuracy of read profiles generated through Bracken. Figure 10 illustrates that 2-step classifiers yield smaller L1 distances, with rspc-based 2-step methods showing the greatest overall reduction. Except for the plant associated samples, which benefit greatly from being classified with the rspc library, there is an increase in L1 distance as the library grows under 1-step classification, which mirrors the loss of specificity in read binning. The 2-step method is able to recover the accuracy loss.

**Fig. 10:**
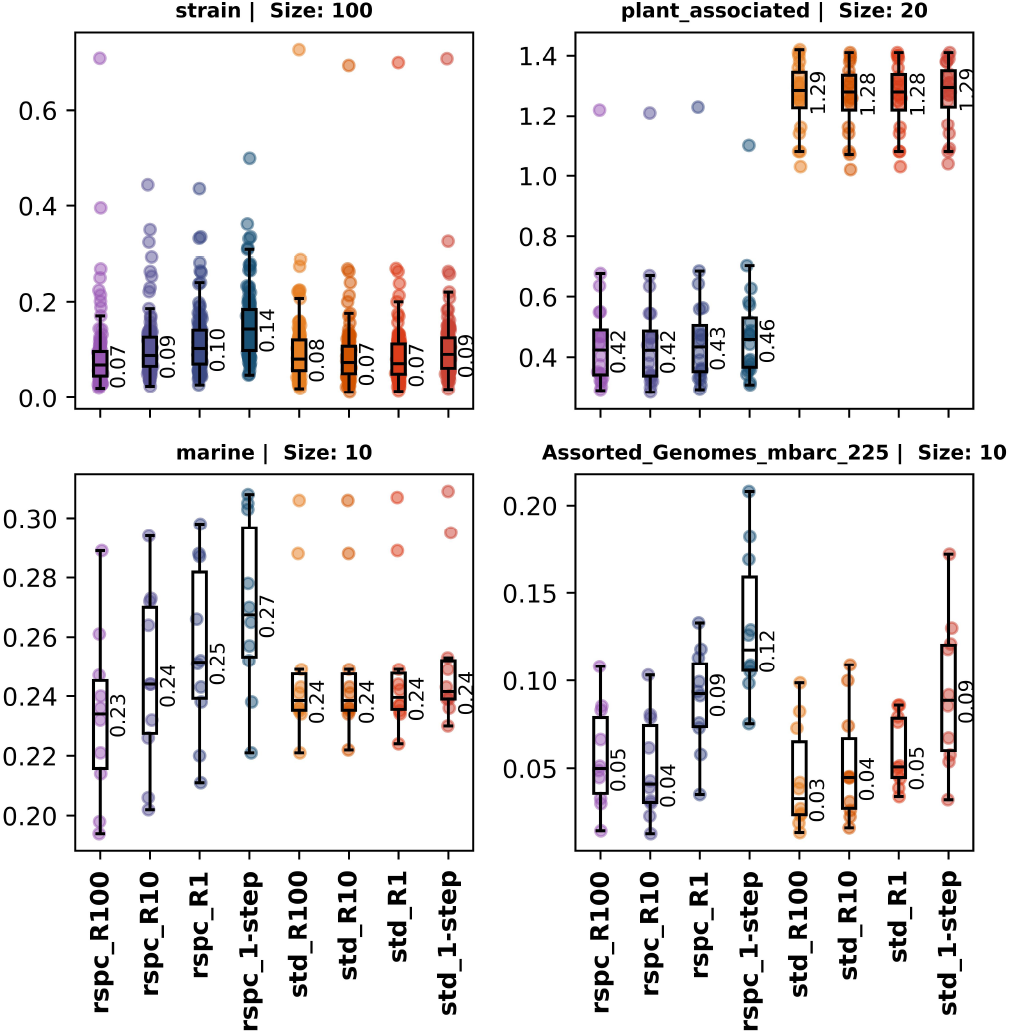
L1 distance, computed between expected read count profile and computed read count profile, for samples in the strain, marine, plant associated and in-silico datasets.

### Gold set classifiers show the potential of 2-step classification with a perfect heuristic

We also analyzed the impact of classification using gold set libraries. The purpose of the 2-step method is to detect as many taxa truly present in the sample as possible, while filtering out taxa not in the sample. The actual set of taxa present in the sample metagenome is referred to as the gold set. To assess the potential maximum performance of the 2-step method, we classified each sample using its respective gold set as a genome filter. Figure 11 presents the performance of the gold set classifications using the rspc and std libraries. It is worth noting that the choice of library is significant, as not all taxa in the gold set necessarily belong to the reference library. Therefore, the final gold set used is the intersection of the ground truth taxa for that sample and the taxon set of the selected reference library.

**Fig. 11:**
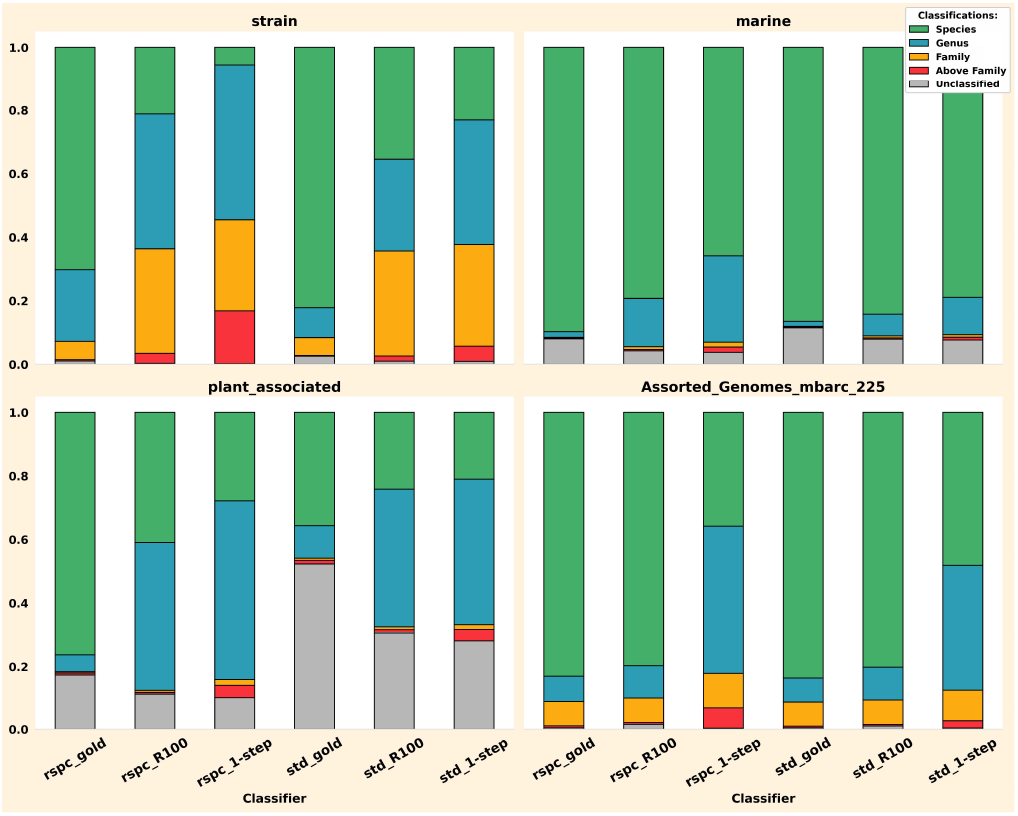
Stacked bar charts to show the performance of various classifiers including rspc and std gold set classifiers. Gold set classifications exhibit a binary tendency, i.e. either a read is classified at a species level or remains unclassified.

We observed that gold set classifiers have a binary tendency: often, a read is either classified precisely at the species level or left unclassified. When dynamic libraries are too tight, they lose the capacity to classify reads from novel genomes as VPs and instead fail to classify them, resulting in FNs. We call this effect ‘squashing the middle’, where VP classifications are converted into either TPs or FNs. Even for samples with reads from real assembled metagenomes like those in the CAMI2 datasets, the gold set performs effectively in this regard. Figure 12 illustrates, somewhat unexpectedly, that the gold set classifications have a very low fraction of FPs, typically about 2 − 5% lower than other classifiers. This is surprising, as our read-based filters generally show a trend of increasing FPs as the library size decreases.

**Fig. 12:**
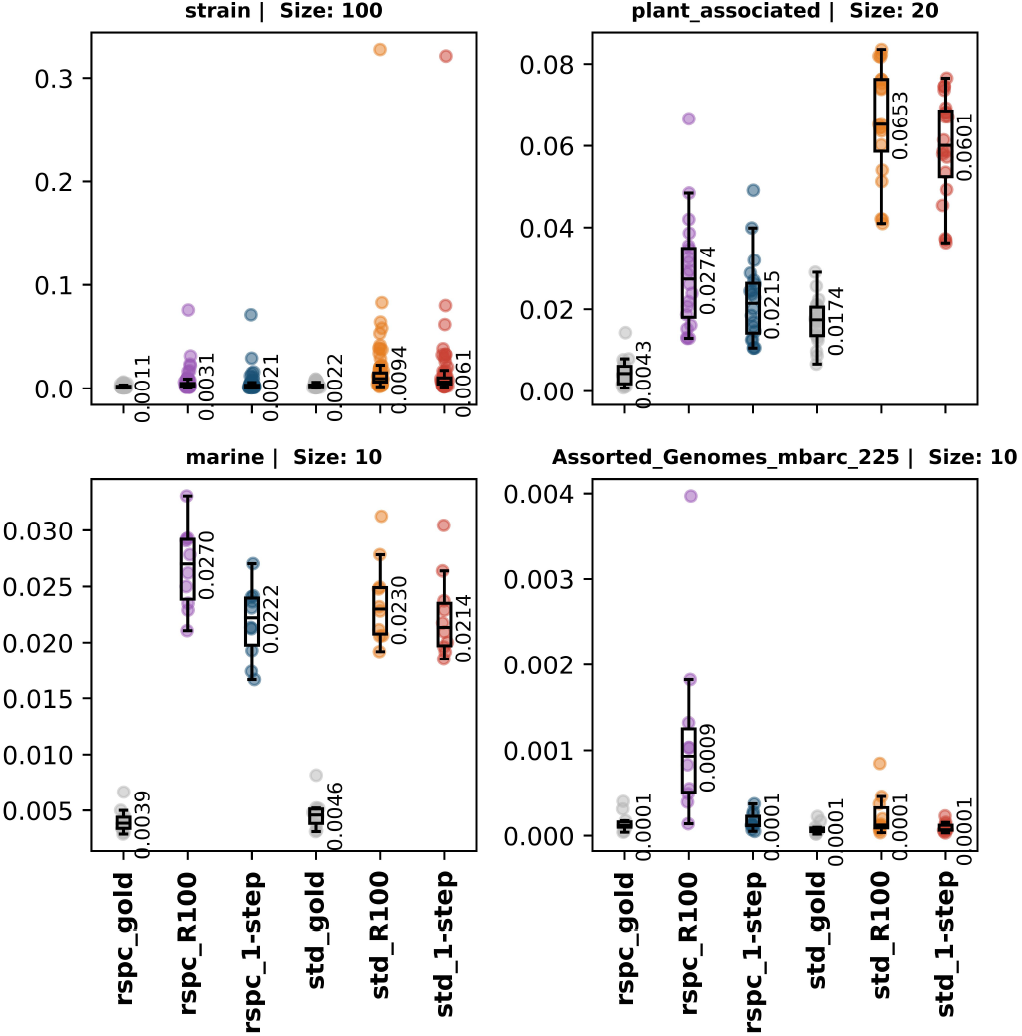
False positive fraction of total sample reads for various 1- and 2-step classifiers as well as gold set classifiers on different datasets. Gold set classifiers tend to have very low percentages of false positives.

The relationship between FPs and library size is not straightforward. Reducing the library size can potentially force reads to be incorrectly classified to the available reference genomes, increasing FPs. However, removing non resemblance genomes also removes these taxa as classification targets, reducing FPs. We speculate that the reason the gold set classifiers have such few FPs is directly due to the maximization of the latter effect.

### The read cutoff heuristic determines taxon set precision

For determining the set of species level taxa found in final output of a classifier, we consider a species as “detected” by following Bracken’s default criterion, which requires at least 10 classified reads for a species to be considered present. For 2-step classifiers with read cutoffs of 10 or higher, the detected species are entirely determined by species-level taxa that pass the heuristic read filter. This follows from the fact that taxa with 10 or more reads in 1-step classification will have at least 10 classified reads in 2-step output.

Figure 13 report taxon set precision and recall for various Slacken classifiers, including theoretical maximum values from gold set classifiers, alongside MetaPhlAn. Unlike read classifiers such as Slacken, which classify all reads, MetaPhlAn estimates taxon abundance by aligning a small fraction of reads to a curated marker database containing unique sequencing regions for different clades. This approach makes it a valuable tool for measuring the relative cellular abundance of taxa in a community. However, while it enables precise taxonomic profiling, it also leads to high precision and low recall when applied to typical microbial community samples.

**Fig. 13:**
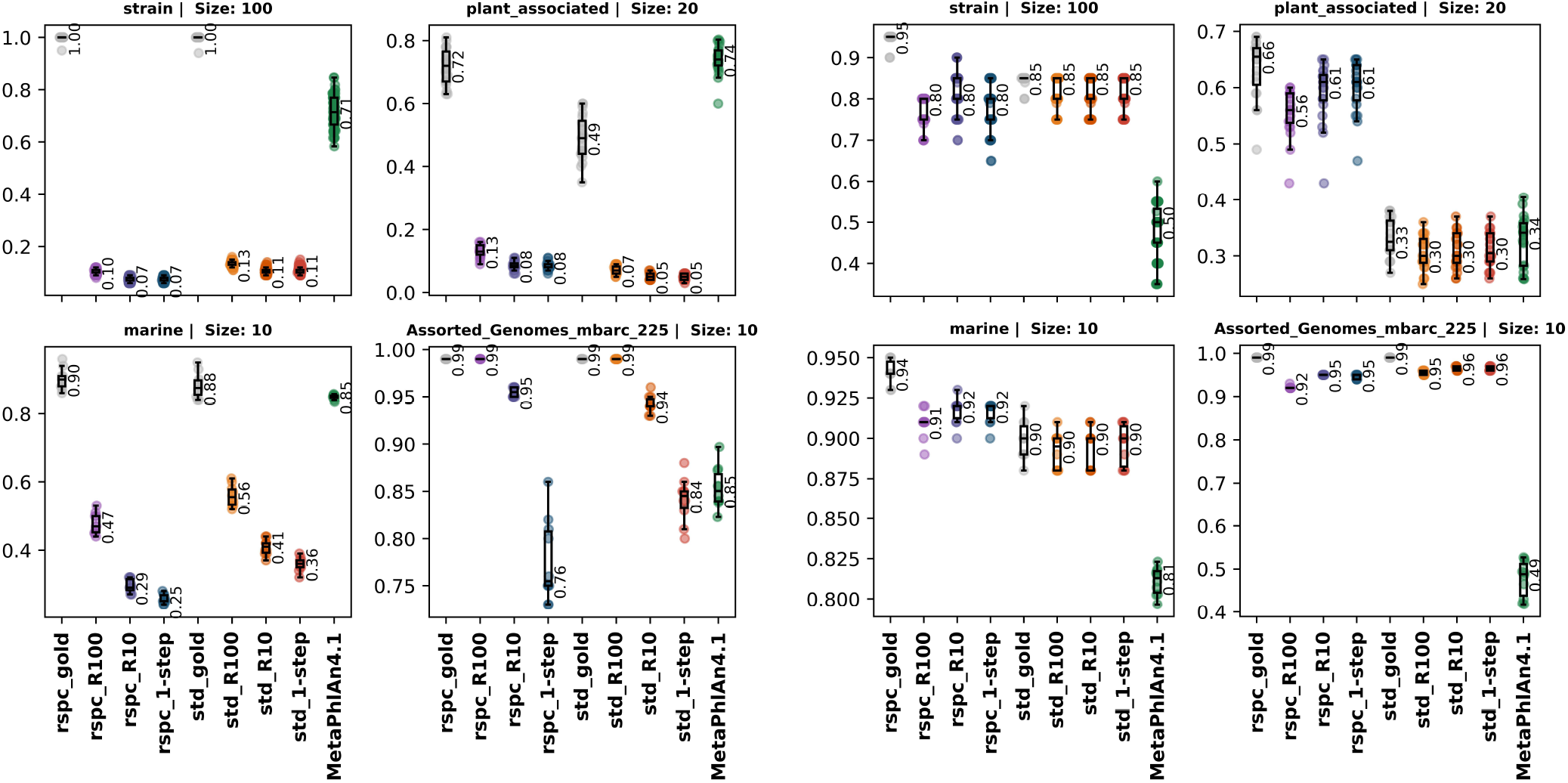
Detected species set precision (left) and recall (right) for each of the four sample families and the various 2- and 1-step classifiers on the *std* and *rspc* library databases as well MetaPhlAn4. A species level taxon is considered ‘detected’ by Slacken if at least 10 reads classify to it in the final output.

For Slacken 2-step, as expected, an increasing read cutoff increases species taxon set precision while reducing recall. Spurious taxa typically have few reads classified, so increasing the read cutoff enhances taxon precision. However, this also removes truly present but low-abundance taxa, as seen in the recall drop from *R*10 to *R*100, highlighting a limitation inherent to the read-based heuristic used. For example, in the case of *R*100 and 10 million reads, this would correspond to the read abundance of 10^−5^ or less for excluded taxa. The gold set classifiers define theoretical performance limits for the 2-step classifiers. MetaPhlAn exhibits a similar pattern but to a greater extent, maintaining very high precision at the cost of significantly reduced recall. Interestingly, despite its marker database (ChocoPhlAn) containing far more taxa from marine and plant datasets than the standard library (see table 2), MetaPhlAn still exhibits poorer recall, underscoring its highly conservative nature.

### 2-step classification with Slacken is fast and efficient

Table 4 breaks down the time and cost requirements of running Slacken 1-step and 2-step classification on 50 samples from the CAMI2 strain madness dataset. Experiments were run using a commercial cloud computing provider. Slacken generally runs faster than Kraken 2 (wall clock time) since it can be distributed on a large number of machines that run in parallel. This does not mean that the number of CPU hours required by Slacken is always smaller. However, because Kraken 2 requires an expensive machine type for large databases (with sufficient RAM), for 2-step classification, the dollar cost per sample was lower for Slacken even when the CPUh requirement was larger. In the case without Bracken, Slacken was nearly four times more efficient than Kraken 2.

**Table 4.**
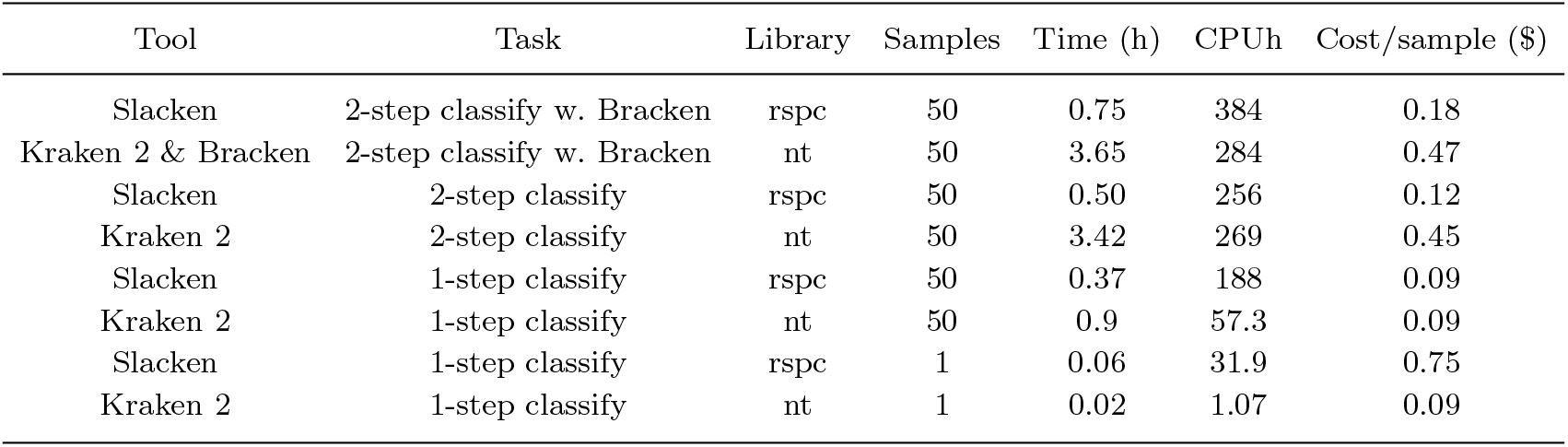
Cost of classify tasks for Slacken and Kraken 2 in a cloud environment.

For 1-step classification, in the case of a single sample, Kraken 2 is more efficient than Slacken, but multisample classification with 50 samples makes the per-sample cost equal for Slacken and Kraken 2. Running the 2-step method with support for Bracken adds about 50% cost per sample due to the complexity of building bracken weights (the Slacken equivalent of `bracken-build`).

## Discussion

More and better reference genomes are constantly becoming available, causing a conflict of aims for classifier tools based on genomic libraries. On the one hand, larger libraries can be preferable since they are good at avoiding false positives and have high sensitivity. On the other hand, a larger library leads to read classifications at higher levels in the taxonomic hierarchy, leading to vagueness, since fewer genomic regions are unique to a single taxon across the whole library. To address this in the context of the Kraken 2 minimizer-LCA classification method, we propose two-step classification. A static, large library is used initially to identify the genomes that are present in samples, and then a second library - a sample-tailored library - is built on the fly from only the genomes actually detected. The second step can then yield species-level classifications with a high specificity while the first step still guards against false positives and retains its sensitivity to a wide range of taxa.

Comparing 1-step classification results from the std library with those from the much larger rspc library, our results confirmed that broad reference databases result in increased vagueness. Using a simple, read-based criterion to include taxa in the dynamic library, we found that the 2-step method effectively mitigated this loss in specificity. The method improves overall assignment quality, shown by a decrease in the sample index between the 1-step and 2-step classifications.

The 2-step method needs an additional parameter over the 1-step method – a filter for selecting genomes in the 2-step library. This significantly impacts classification. If this filter is too permissive, specificity is reduced, making classifications similar to those of the 1-step method (or Kraken 2). On the other hand, an overly restrictive filter risks excluding genomes that should be represented in the sample. Incorrectly excluding a genome forces any reads from that genome to appear either as false negatives or as false positives from some other genome. This effect is amplified by the smaller size of the 2-step library, which can lead to notable decreases in minimizer-assigned LCAs.

From a quantitative biology perspective, applying a taxon read cutoff is practical, as kit and laboratory contaminants often introduce trace-level species, making ultra-low abundance taxa potential contaminants [17]. While this can critically impact metagenomics in low biomass communities [9], sensitive classification tools (like Slacken and Kraken) can incorrectly detect contaminant taxa from even a single sequenced read, leading to many low-abundance false positive taxa in high biomass communities. Such taxa are typically addressed post-classification, where the classifier output is filtered for species above a certain read (or taxonomic) abundance threshold [18]. However, this could lead to removal of low abundance taxa that are actually present. By eliminating ultra low abundance (*R* = 1, 10, 100) taxa prior to the 2-step classification, we are able to buy back precision, potentially increasing reads classified to true low abundance taxa in the final output.

The R100 heuristic performed well as a genome filter throughout. 2-step R100, using the rspc library, greatly increased species level classifications compared with 1-step classification. Every sample family that we studied showed improvement. For the CAMI2 strain samples, while using the rspc genome library, the fraction of species-level classifications was increased by 3.5x relative to 1-step, while for in silico samples (Assorted genomes mbarc 225), they were increased by 2.2x to a total of 80%. Classifications at higher levels (above family) were shifted downward substantially: from 17% to 3% in the strain samples, and from 7% to 1% in the in silico samples. The fractions of false positives and false negatives changed only very slightly. Based on our experiments, we believe that using the R100 heuristic together with a maximally large initial library would be a good starting point for most users. For users who are concerned about R100 possibly missing some taxa, the R10 heuristic is a more conservative alternative.

Using a read-based cutoff to reclassify reads has similarities with Bracken’s default mode of operation. Bracken also employs a cutoff (by default 10) to select target species for redistribution of reads. However, Bracken redistributes hypothetical reads to arrive at an estimate of the true number of reads for each taxon, whereas our 2-step approach assigns specific reads to taxa.

Using ground truth taxon sets (gold set classifiers), we demonstrate the best performance achievable by a perfect genome filter. Although we found that the R100 filter performed well on most metrics, there is a large gap between R100 performance and gold set classifier performance that remains unexplored. Future investigations could explore more stringent read filters (higher *R* cutoffs) or other types of taxon selection criteria. For example, KrakenUniq [19] extends Kraken and Kraken 2 by tracking the number of distinct k-mers detected per taxon, a signal that could also be used as part of a 2-step selection filter.

While our CAMI2 results do not indicate significant problems with FP and FN reads relative to 1-step classification, unfortunately, removing taxa not detected by the read cutoff heuristic carries the risk of missing novel or unseen genomes. For example, if some reads classify at genus level but no species level descendants satisfy the cutoff under that genus, the entire genus would be excluded from the second library. We are actively investigating how to best handle this in the context of a 2-step method.

Tuning the read cutoff parameter may involve multiple conflicting goals. For example, we observed that a lower cutoff can increase taxon set recall. On the other hand, a higher cutoff favors taxon set precision as well as read profile accuracy (lowering the read index overall, increasing specificity). In addition, factors such as sequencing depth of samples may directly affect the choice of this value: for a 10x greater sequencing depth, we would expect a 10x higher R value to yield a comparable result. We hope to establish clear guidelines in future work. Currently, we recommend users to explore parameters such as sample grouping and the read threshold and compare Slacken’s outputs with either a known ground truth or results from more thorough method, such as sequence alignment. This problem is not new to Slacken: when using Kraken 2 and Bracken, parameters such as confidence threshold and a read cutoff must be chosen, and here too the choice of parameters must somehow be validated.

When using Slacken, we found that the cost and runtime per sample of the 2-step approach is comparable to regular Kraken 2 classification when using large sample libraries and when classifying multiple samples together. However, multisample classification affects read assignments as well as cost. We classify related samples together, since every sample can, through the genome detection heuristic, influence the 2-step library and thus the second step classification. We believe that as long as samples carry information about the same signal (for example, if they originated in the same environment), grouping more samples together should always help accuracy, since more faintly present taxa might cross the initial read count threshold and be correctly picked up by the first step. On the other hand, grouping unrelated samples together could reduce specificity.

Unfortunately, multisample classification could make reproducibility difficult in a situation where sample sets grow or change over time. We are investigating best practices for such a scenario as part of our ongoing work. In some cases, Bracken may be able to mitigate reproducibility problems by compensating for any changes in Slacken’s second step taxon set. Alternatively, the two-step method can be used on a single sample at a time. This is always reproducible, although more costly.

There are generally two approaches to analyzing a sample metagenome: one aims to classify as many reads as possible, even at higher taxonomic ranks (especially for novel taxa), to capture a broad view of the entire sample’s taxonomic diversity, despite potential novel genomes being present. The other focuses on precisely classifying reads from taxa already represented in the reference library, thereby limiting classification to known organisms. Slacken’s 1-step (or Kraken 2) approach aligns with the first method when using large, representative libraries containing genomes spanning the tree of life. However, the second approach—achieving precise classification with a focus on known taxa—has been challenging because it traditionally requires smaller libraries. Using reference libraries from specific branches of the taxonomic tree has been shown to skew results, potentially increasing false positive classifications due to shared genomic regions and library contamination [7]. Slacken’s 2-step method addresses this issue by enhancing classification specificity while preserving the robustness of using a broad, comprehensive reference.

For taxon set detection, Kraken 2 and MetaPhlAn also exemplify two contrasting approaches-one maximizing taxon recall and the other prioritising precision. Each serves a distinct purpose: Kraken 2 for read classification and MetaPhlAn for taxon profiling. We believe that by tuning parameters such as the read cutoff, Slacken’s 2-step method could be tuned to either of these purposes. Slacken’s heuristic cutoff quantitatively determines the detected taxon set. This presents an opportunity for future work to incorporate additional information, such as marker databases or direct MetaPhlAn output, to further refine 2-step library taxon selection and reduce spurious detections. Additionally, alternative heuristics could be designed to make Slacken more adaptable, allowing it to span the spectrum between Kraken’s recall-focused approach and MetaPhlAn’s precision-oriented profiling. Exploring such heuristics could enable classifier behaviour to be tailored to specific problem contexts.

Beyond 2-step classification, there are other potential ways to improve the k-mer/minimizer approach to classifying sequences. For example, longer minimizers or k-mers could capture more information from each genome, especially in regions that are not conserved between multiple genomes. In this work we have mainly focussed on 2-step classification with the case of the widely accepted Kraken 2 standard parameters, which are 31-minimizers of 35-mers. In future work we hope to also explore longer minimizers for both 1-step and 2-step classification.

## Conclusion

K-mer based metagenomic binning and profiling continues to be widely used. However, lowest common ancestor (LCA) based algorithms do not scale well as libraries keep growing, since classifications become increasingly vague when LCA taxa move higher up in the taxonomic tree. We find that sample-tailored minimizer libraries, which filter out non-resemblance genomes in a second step prior to classification based on a simple read count heuristic, overcome this limit by allowing the initial library to scale freely while the tailored library can produce specific classification results. This 2-step method moves classifications lower in the tree of life across all levels in the taxonomic hierarchy, a result that we observed for both the sequenced CAMI2 samples and *in silico* samples. Furthermore, 2-step classification with Slacken is comparable in cost to regular Kraken 2 classification when classifying multiple samples with a large library. Sample-tailored libraries set a new standard for precision and allow k-mer-based LCA methods to achieve better performance as genomic reference libraries keep growing.

## Source code and data availability

Slacken is available as open source software under the GPL license at https://github.com/JNP-Solutions/Slacken. The same repository also includes the tools and scripts used for constructing the std and rspc libraries, and for sample filtering and evaluation of classifiers. We have applied for AWS open data sponsorship for the pre-built std and rspc libraries. In the meanwhile, we are happy to provide them on a need basis.

Supplementary data is available at https://github.com/JNP-Solutions/Slacken/tree/master/metrics.

## Author contributions statement

J.N.P and N.B conceived of the study, conducted experiments and analysed data. J.N.P. developed the Slacken software. J.N.P, N.B and S.G wrote and reviewed the manuscript.

## Acknowledgments

This work was partially supported by the ONRG Grant for the Nobel Turing challenge to The Systems Biology Institute (Grant number: N62909–21–1–2032) for J.N.P, N.B and S.G.

