## Supplementary Figures for "Precise and scalable metagenomic profiling with sample-tailored minimizer libraries"

Table 4: Fraction of total reads classified at different ranks for each dataset and classifier. A.G. is short for assorted genomes.

| Dataset | Group | rspc_1-step | rspc_R1 | rspc_R10 | rspc_R100 | std_1-step | std_R1 | std_R10 | std_R100 |
|---|---|---|---|---|---|---|---|---|---|
| A.G._mbarc_225 | Species | 0.36 | 0.47 | 0.74 | 0.80 | 0.48 | 0.60 | 0.74 | 0.80 |
| | Genus | 0.46 | 0.41 | 0.16 | 0.10 | 0.39 | 0.29 | 0.17 | 0.10 |
| | Family | 0.11 | 0.10 | 0.08 | 0.08 | 0.10 | 0.09 | 0.08 | 0.08 |
| | Above-Family | 0.07 | 0.01 | 0.01 | 0.01 | 0.02 | 0.01 | 0.00 | 0.00 |
| | Unclassified | 0.00 | 0.00 | 0.01 | 0.02 | 0.00 | 0.00 | 0.01 | 0.01 |
| marine | Species | 0.66 | 0.69 | 0.73 | 0.79 | 0.79 | 0.81 | 0.82 | 0.84 |
| | Genus | 0.27 | 0.25 | 0.21 | 0.15 | 0.12 | 0.10 | 0.09 | 0.07 |
| | Family | 0.02 | 0.01 | 0.01 | 0.01 | 0.01 | 0.01 | 0.01 | 0.01 |
| | Above-Family | 0.02 | 0.01 | 0.01 | 0.00 | 0.01 | 0.00 | 0.00 | 0.00 |
| | Unclassified | 0.04 | 0.04 | 0.04 | 0.04 | 0.08 | 0.08 | 0.08 | 0.08 |
| plant_associated | Species | 0.28 | 0.32 | 0.36 | 0.41 | 0.21 | 0.22 | 0.23 | 0.24 |
| | Genus | 0.56 | 0.55 | 0.51 | 0.47 | 0.46 | 0.46 | 0.44 | 0.43 |
| | Family | 0.02 | 0.01 | 0.01 | 0.01 | 0.02 | 0.01 | 0.01 | 0.01 |
| | Above-Family | 0.04 | 0.01 | 0.01 | 0.01 | 0.04 | 0.02 | 0.01 | 0.01 |
| | Unclassified | 0.10 | 0.11 | 0.11 | 0.11 | 0.28 | 0.29 | 0.30 | 0.31 |
| strain | Species | 0.06 | 0.07 | 0.13 | 0.21 | 0.23 | 0.28 | 0.29 | 0.36 |
| | Genus | 0.49 | 0.53 | 0.50 | 0.42 | 0.39 | 0.35 | 0.35 | 0.29 |
| | Family | 0.29 | 0.33 | 0.33 | 0.33 | 0.32 | 0.33 | 0.33 | 0.33 |
| | Above-Family | 0.17 | 0.07 | 0.04 | 0.03 | 0.05 | 0.03 | 0.02 | 0.02 |
| | Unclassified | 0.00 | 0.00 | 0.00 | 0.00 | 0.01 | 0.01 | 0.01 | 0.01 |

### 1 Classification rank distribution

Table 4 gives the distribution of read ranks for each dataset and classifier.

### 2 Visualisation of the supplementary tables S1 & S2

A collection of graphical visualisations of selected data from tables S1 and S2 with various classification metrics with 11 different classifiers and 6 datasets, using a confidence of 0.15.

- Figure 1 : True Positive read classifications.
- Figure 2 : False Positive read classifications.
- Figure 3 : Vague Positive read classifications.
- Figure 4 : False Negative read classifications.
- Figure 5 : Sample Index boxplot
- Figure 6 : Sample soft-index boxplot
- Figure 7 : L1 (Manhattan) distance from ground truth.

- Figure 8 : LSE (Euclidean) distance from ground truth.

### 3 Index metrics

Properties of the index and s-index:

1. The s-index can be used to cross-compare classifiers that use different representative genome libraries. Figure 6 shows the usefulness of the s-index metric. We know that larger genome libraries can allow us to classify more reads but also tend to blunt read classifications[1], both tending to increase the s-index[2]. It is clear that if the index of an rspc based classifier is lower than an std based classifier (for a given sample), then the quality of read classifications can be considered to be superior. I.e. we expect more reads to be classified at smaller per-read indices.

2. Note that the above is not necessarily true for the index, as simply classifying FN labelled reads, even at a very high rank, will tend to lower the index. This means that read classifications can become very blunt as long as many unclassified reads are classified.

3. The index can be used to internally-compare classifiers that use the same representative genome library. Figure 5 shows the usefulness of the index. We know that shrinking a genome library (as the 2-step method does) sharpens read classifications at the cost of generating more FN read classifications. The index tends to increase with increase in FNs. It is clear that if the index of a classifier with a higher read cutoff (smaller 2-step library) is lower than that of a classifier with a lower read cutoff, using the same representative genome library, then the quality of read classifications can be considered to be superior.

4. Note that this is not necessarily true for the s-index, as an increase in FN reads tends to decrease this metric. It is possible then that read classifications are not any sharper, we're simply classifying fewer reads.

5. Finally, the index can be used to cross-compare classifiers on different representative genome libraries, as a cautionary upper bound on the quality of 2-step classifications using large libraries. The index tends to decrease when going from an std to an rspc based classifier simply due to more reads being classified.

[1] This is not a strict rule. It possible, however remote, that large libraries introduce new minimizers that contribute to sharpening classifications of already classified reads. This would mean that an already classified read has minimizer that didn't exist in the smaller library. This is generally very unlikely.

[2] Note that blunting of read classifications leads to a higher per-read index and classifying *fn* labelled reads will lead to a change in index from 0 to the distance between the classified taxon and the ground truth

Figure 5 shows how the index captures the strength of a taxonomic read binner. Note that across representative genome libraries, a lower index could correspond to more accurate classifications or a greater number of reads classified or both. For classifiers using the same 1-step index, a lower index directly indicates more accurate classifications and a lowering (on average) of the individual index values for vague positive read classifications.

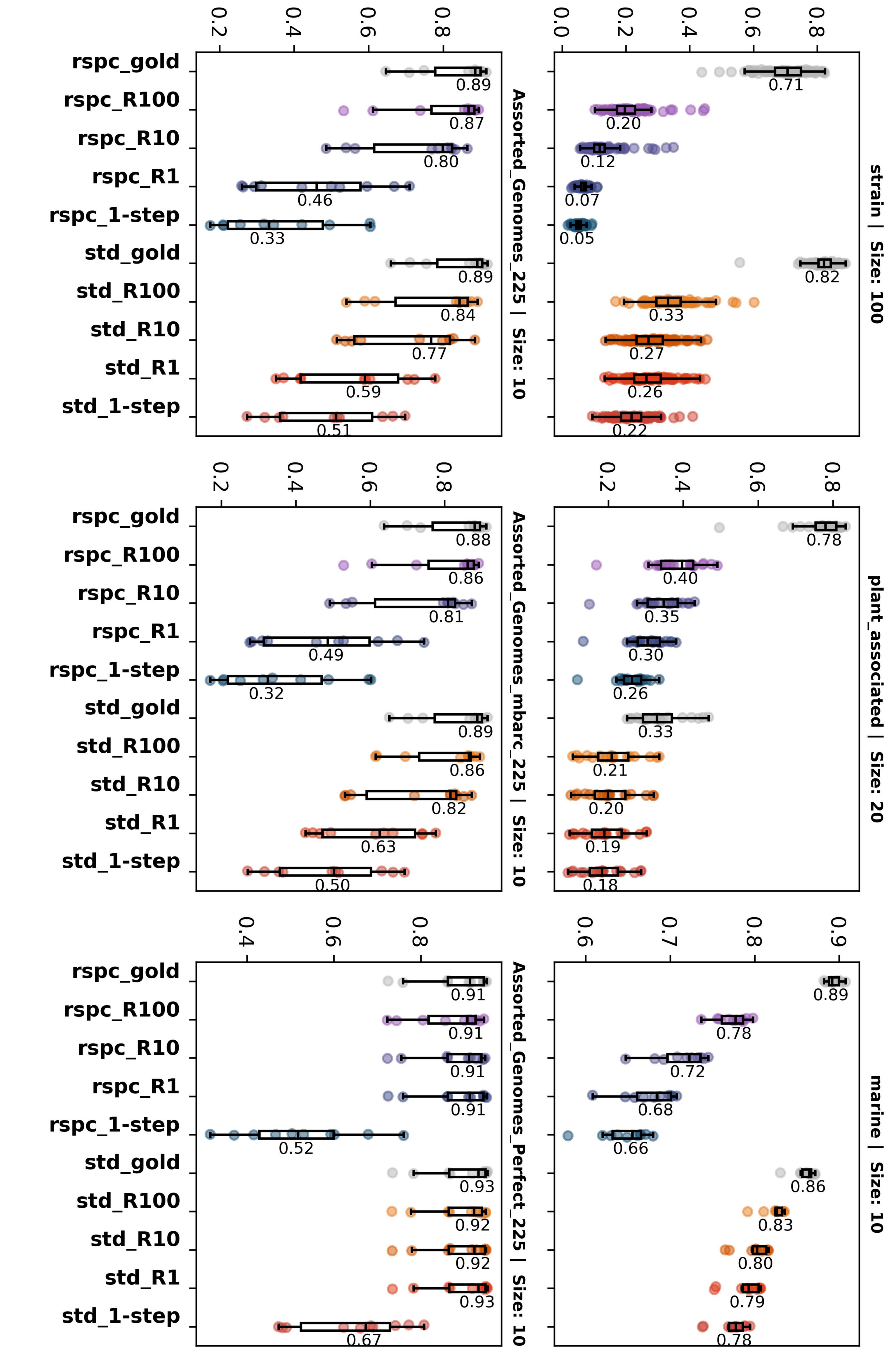


Figure 1: (Species Level) True positive fraction of total sample reads for various 1- and 2-step classifiers as well as gold set classifiers on different datasets.

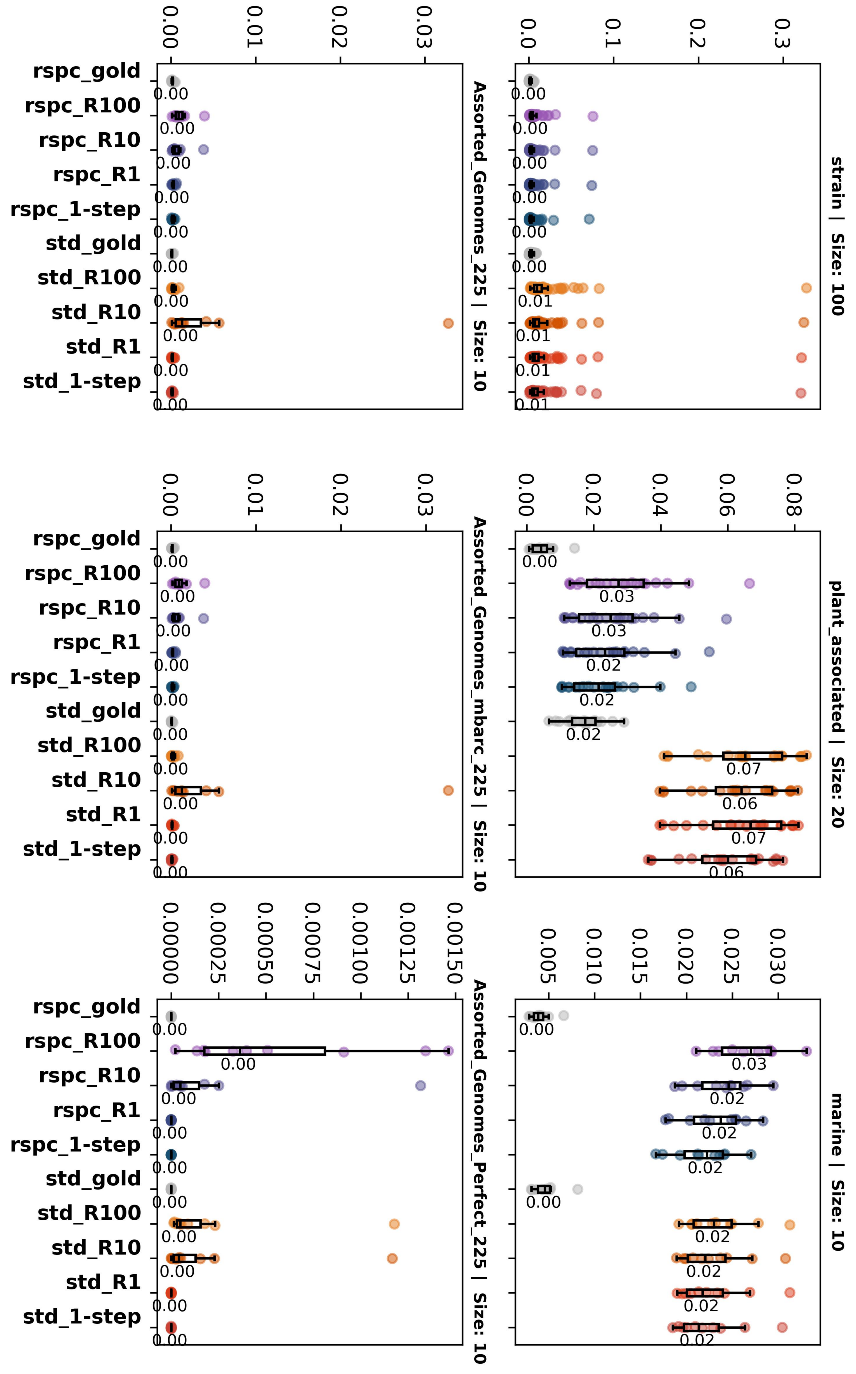


Figure 2: (Species Level) False positive fraction of total sample reads for various 1- and 2-step classifiers as well as gold set classifiers on different datasets.

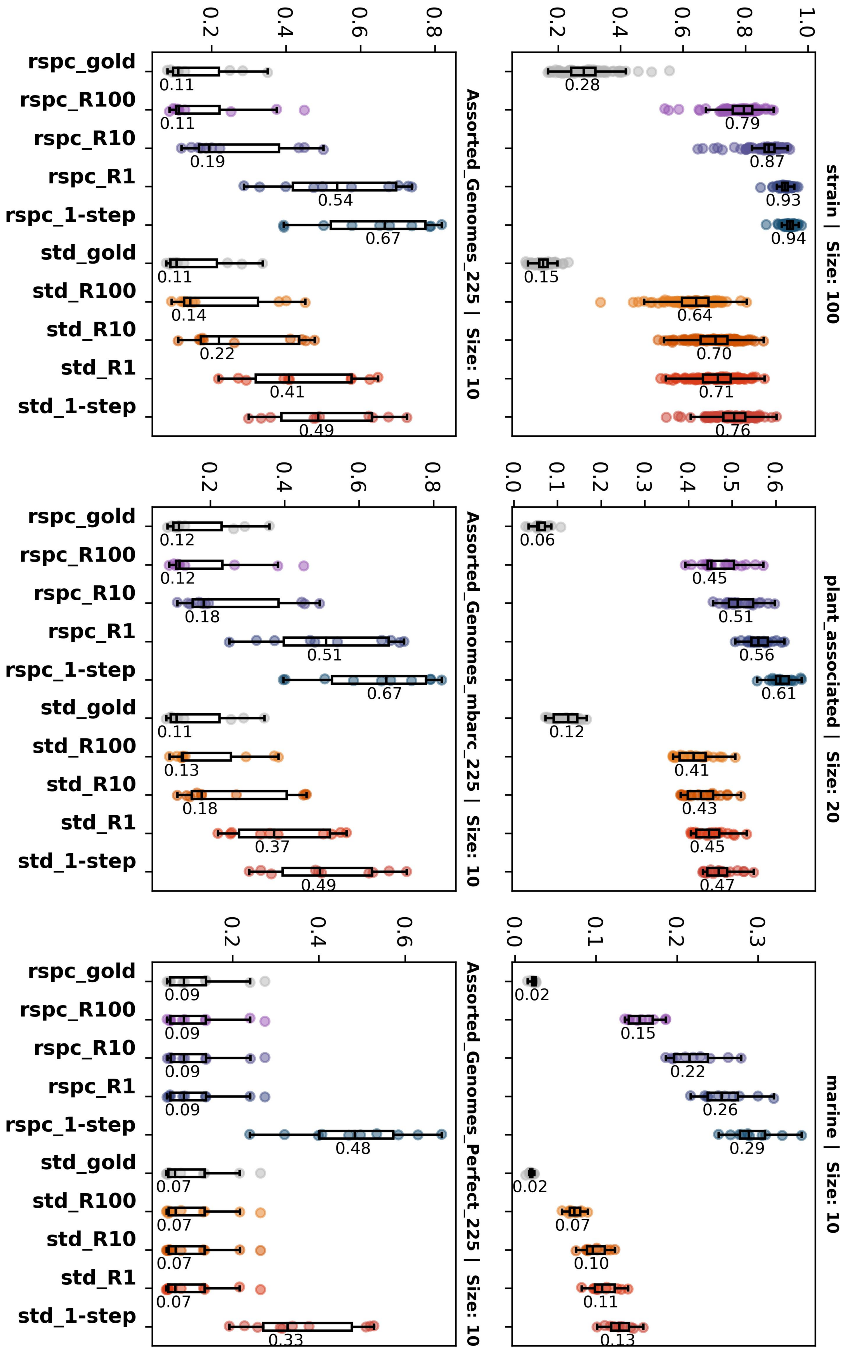


Figure 3: (Species Level) Vague positive fraction of total sample reads for various 1- and 2-step classifiers as well as gold set classifiers on different datasets.

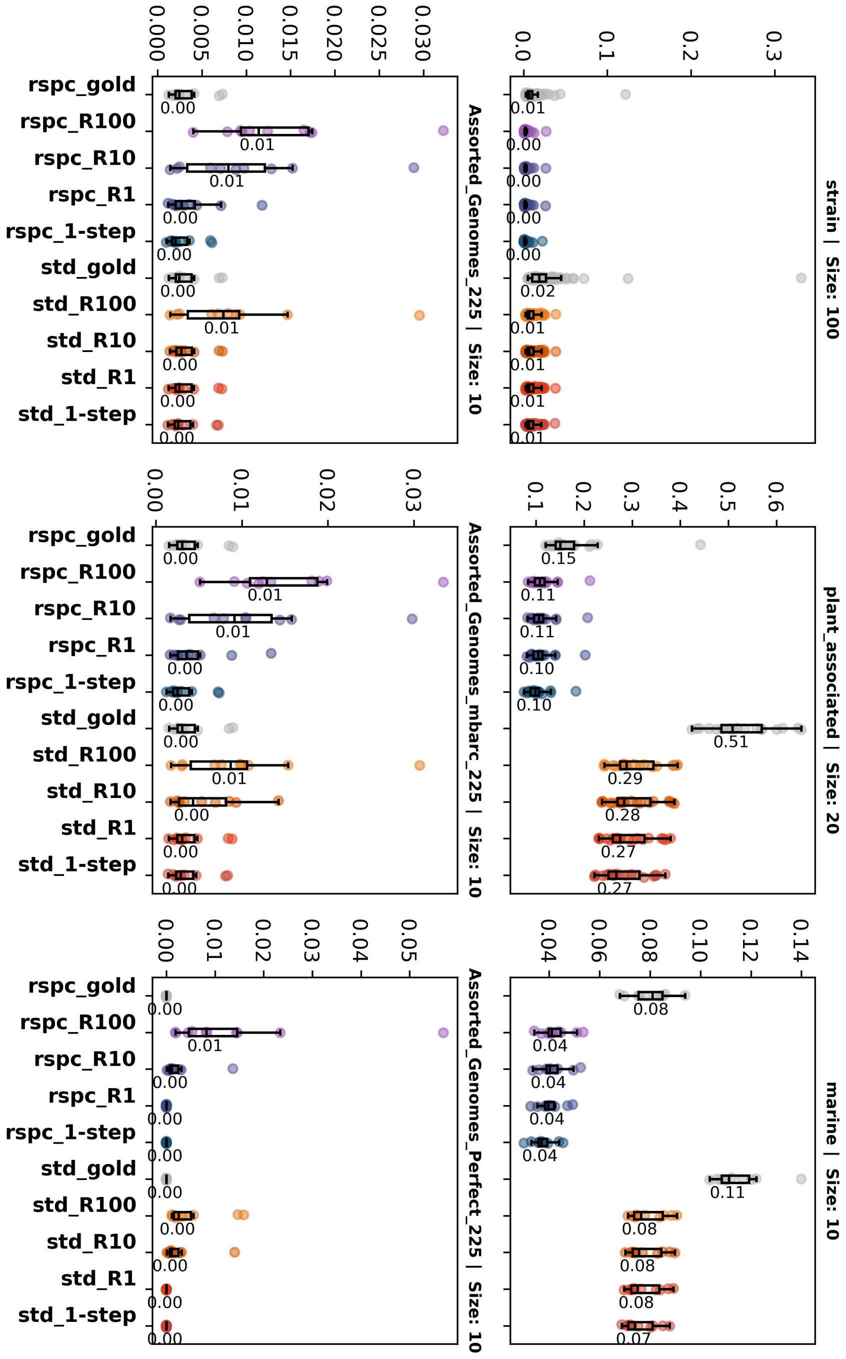


Figure 4: (Species Level) False negative fraction of total sample reads for various 1- and 2-step classifiers as well as gold set classifiers on different datasets.

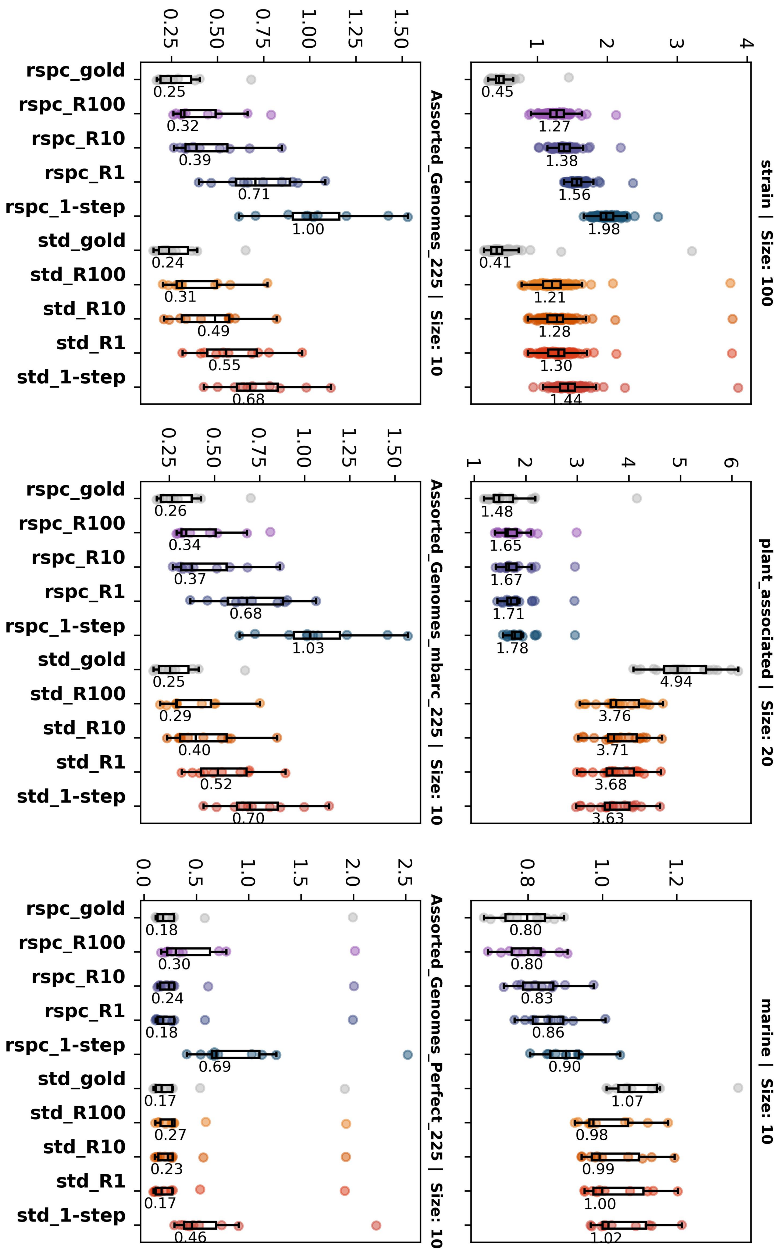


Figure 5: (Species Level) Boxplots of sample index values for strain, plant_associated, marine and mbarc datasets with various 1- and 2-step classifiers. Sample index is computed by taking a weighted average of all sample reads, giving a weight of 0 to TPs and 9 to FPs and FNs. VP reads are given a weight corresponding to the number of ranks between the true taxon and the classfied taxon label for that read.

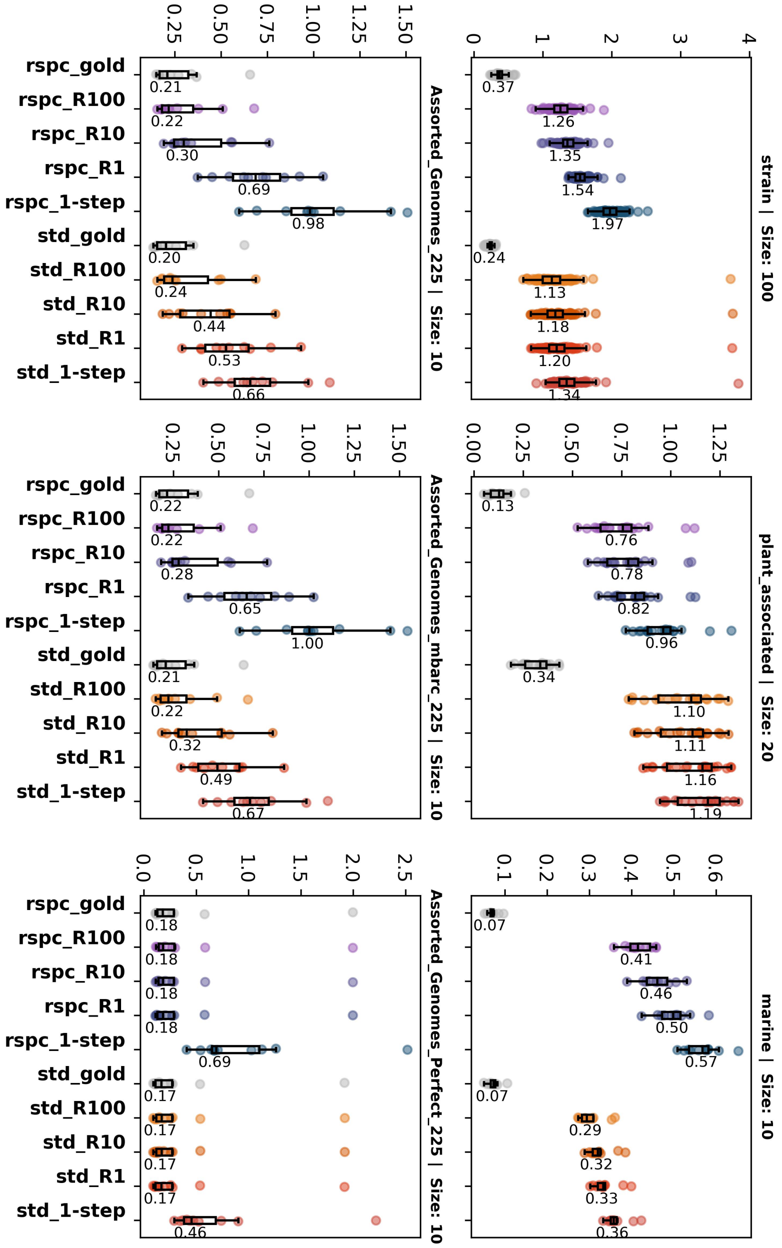


Figure 6: (Species Level) Boxplots of soft-index values for strain, plant_associated, marine and mbarc datasets with various 1- and 2-step classifiers. Soft-index is computed by taking a weighted average of all sample reads, giving a weight of 0 to TPs and FNs and 9 to FPs. VP reads are given a weight corresponding to the number of ranks between the true taxon and the classfied taxon label for that read.

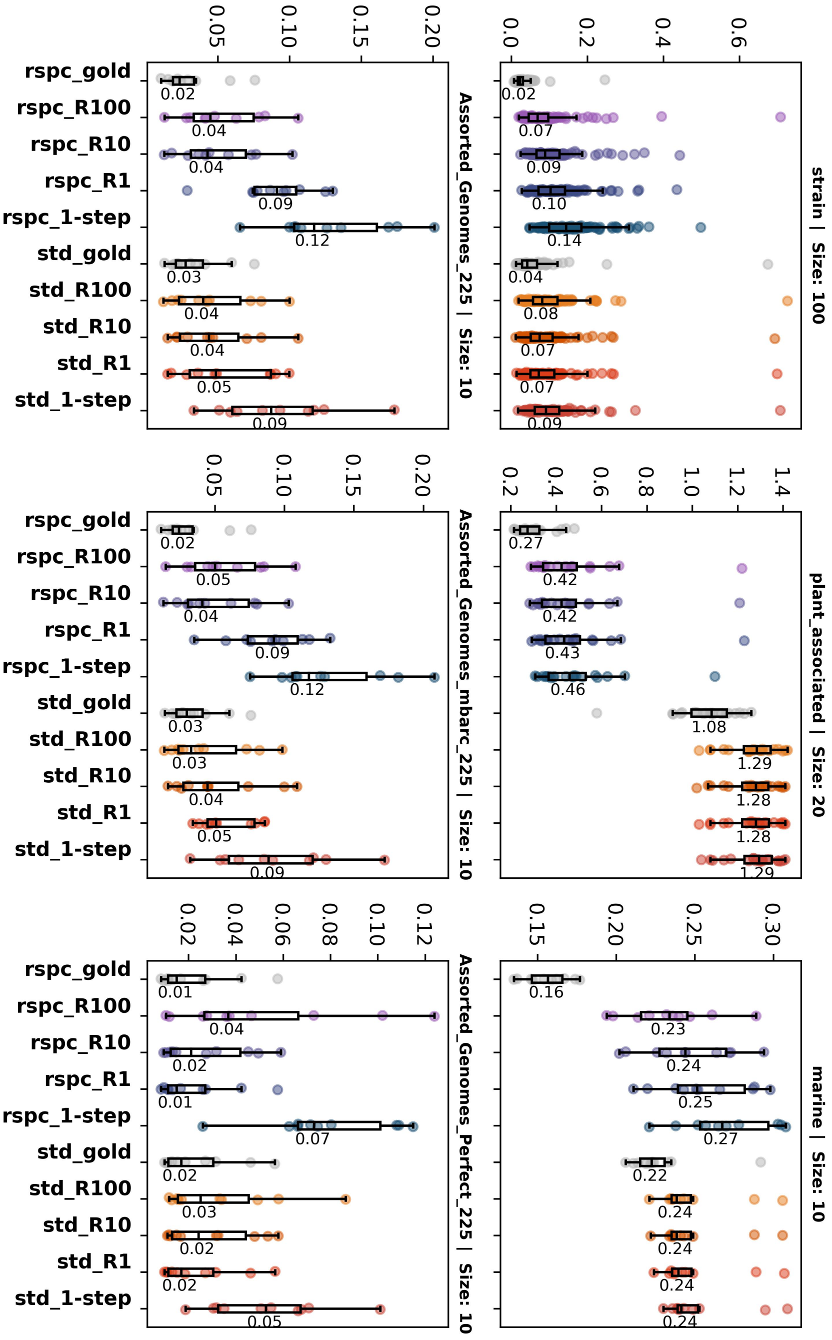


Figure 7: L1 distance, computed between expected read count profile and computed read count profile, for samples in the strain, marine, plant_associated and in-silico datasets.

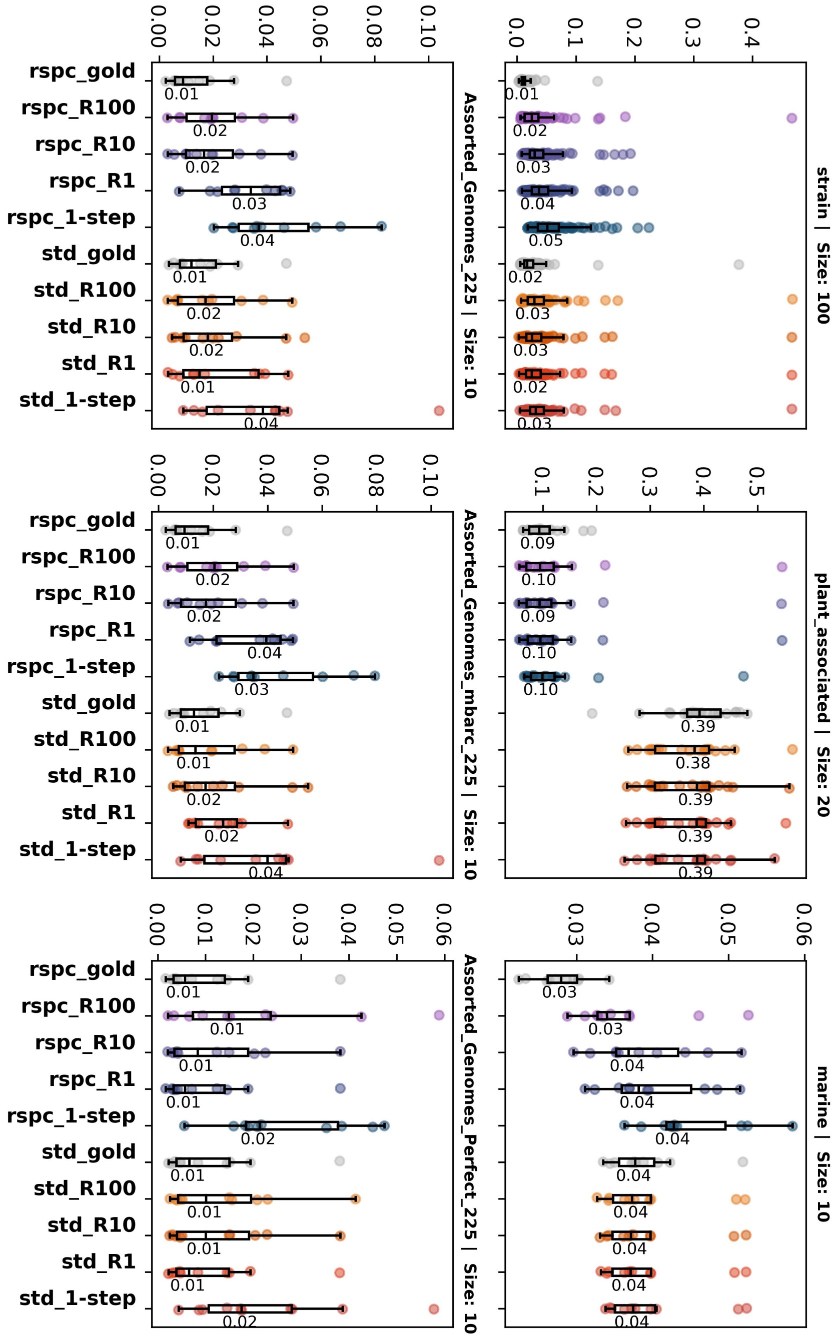


Figure 8: LSE distance, computed between expected read count profile and computed read count profile, for samples in the strain, marine, plant_associated and in-silico datasets.
